# Temporal and spatial frameworks supporting plant responses to vegetation proximity

**DOI:** 10.1101/2022.06.16.496398

**Authors:** Pedro Pastor-Andreu, Jordi Moreno-Romero, Mikel Urdin-Bravo, Julia Palau-Rodriguez, Sandi Paulisic, Elizabeth Kastanaki, Vicente Vives-Peris, Aurelio Gomez-Cadenas, Antía Rodríguez-Villalón, Jaime F. Martínez-García

## Abstract

After perception of vegetation proximity by the phytochrome photoreceptors, shade-avoider plants initiate a set of responses known as the Shade Avoidance Syndrome (SAS). The shade-induced de-repression of active phytochrome B (phyB) releases the repression imposed over the PHYTOCHROME INTERACTING FACTORs (PIFs). In *Arabidopsis thaliana* seedlings, this mechanism triggers rapid and massive changes in gene expression, increases auxin production in a SHADE AVOIDANCE 3-dependent manner and promotes hypocotyl elongation. Other components, such as phyA and ELONGATED HYPOCOTYL 5 (HY5), also participate in the shade regulation of the hypocotyl elongation response repressing it. However, it is less clear how phyA and HY5 interact with PIFs to regulate this response. Our physiological, genetic, cell biology and transcriptomic analyses showed that these components are organized in two main branches, and incorporate into the model for the regulation of shade-induced hypocotyl elongation the temporal and spatial functional importance of the various SAS regulators analyzed in here. They also indicated that PIFs and HY5, belonging to separate branches, target common genes whose expression is rapidly modulated by shade. This transcriptional relationship, however, changes after longer shade-treatments, suggesting that it is a dynamic convergence point to modulate the hypocotyl elongation.

## INTRODUCTION

When plants grow in high density, the close proximity of vegetation might obstruct sunlight. As a consequence, light can become a limited resource and pose a threat for plant survival. Therefore, plants have adopted contrasting avoidance or tolerance strategies to deal with vegetation proximity or shade. Specifically, when shade-avoider (sun-loving) plants face this scenario, they display a set of responses known as the shade avoidance syndrome (SAS). Some of the SAS responses aim to acclimate photosynthesis in a situation of light shortage caused by the presence of neighboring plants; others focus on redirecting growth to escape from shade by promoting either stem elongation and/or apical dominance (reduced branching), or flowering to produce seeds and ensure the species survival [1–3]. At the seedling stage, hypocotyl elongation is likely the best characterized and most conspicuous SAS response in *Arabidopsis thaliana* [2, 4] and the focus of this manuscript.

Plants detect neighbor vegetation as changes in the red (R) to far-red light (FR) ratio (R:FR), a parameter that is strongly affected by the proximity of vegetation. Under low planting density, the direct sunlight changes the intensity during the day but the R:FR (> 1.2) remains relatively constant [5]. By contrast, under higher planting density two different scenarios can be found: plant proximity or canopy shade. When neighboring plants do not shade each other but are close enough, they reflect mainly FR from sunlight compared to other wavelengths. The reflected FR combines with sunlight and results in a moderate decrease in the R:FR (R:FR 0.5-0.3) without reducing light intensity, a signal informing about plant proximity. When neighboring vegetation is denser forming a plant canopy, photosynthetic pigments of the upper leaves act as selective filters that preferentially absorb and deplete blue and R from sunlight, but transmit most FR. Plant canopy shade presents a drastic reduction of R compared to the FR that results in a very low R:FR ratio (R:FR < 0.06) and a low light intensity in the photosynthetic active radiation region [4, 6, 7]. Importantly, in both cases, the reduced R:FR acts as a reliable signal indicative of the nearby presence of vegetation that is perceived by the phytochrome photoreceptors (Figure 1). Hypocotyl (stem) elongation is the strategy to overgrow neighboring seedlings to reach better (non blue- and R-deprived) light conditions.

**Figure 1.**
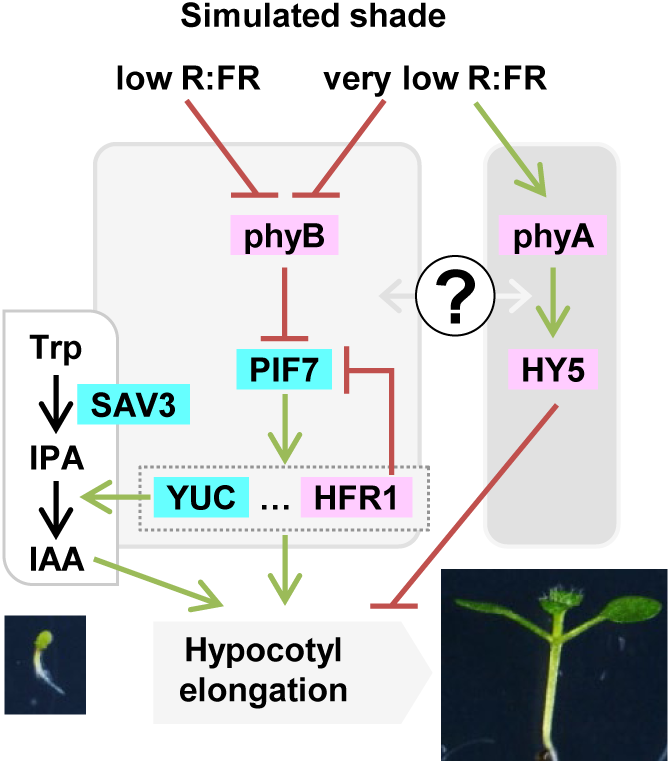
Simplified model that depicts the genetic components analyzed in this work involved in plant neighbor detection. Color indicates the positive (blue) or negative (pink) contribution to the shade-induced hypocotyl elongation. Low and very low R:FR inactivates phyB, which in turn releases the repression imposed on PIFs, mainly PIF7. PIFs subsequently activate the expression of dozens of genes, such as *YUC* genes, involved in IAA production in concert with *SAV3*. The rapid and transient IAA production is required for the hypocotyl elongation. PIF7 also promotes the expression of *HFR1*, that inhibits PIF activity. Under very low R:FR, phyA accumulates to sufficient levels to inhibit hypocotyl elongation, an effect that seems to involve changes in *HY5* expression and/or activity. Overall, exposure to low and very low R:FR activates a transcriptional network that induces shade avoidance hypocotyl response.

As molecular switches, phytochromes exist in two photoconvertible isoforms (an inactive R-absorbing Pr form and an active FR-absorbing Pfr form) that are present in an equilibrium that depends on the prevailing R:FR. Under high R:FR (low vegetation density), most phytochromes are in the active Pfr forms and SAS is suppressed, whereas under low R:FR (high vegetation density), the photoequilibrium moves toward the inactive Pr form and SAS is induced. From the five phytochromes characterized in *A. thaliana* (phyA to phyE), phyA and phyB have the main roles in controlling SAS responses. Genetic and physiological analyses indicate that photostable phyB is the major phytochrome controlling the SAS and other photostable phys act redundantly with phyB in the control of some SAS responses [2, 4]. Additional genetic analyses also showed that phyA, the only photolabile phytochrome, has an antagonistic role over phyB in the SAS control, particularly under very low R:FR, that mimics plant canopy shade. Under low R:FR, wild-type seedlings present a similar hypocotyl elongation as *phyA* whereas *phyB* displays a higher elongation. In contrast, under very low R:FR, wild-type and *phyB* seedlings elongate less than when grown in low R:FR and *phyA* seedlings present an exaggerated hypocotyl length. This indicates that phyB is deactivated by both, proximity (low R:FR) and canopy shade (very low R:FR) whereas phyA activity is induced only by very low R:FR (Figure 1) [4, 8, 9]. Consistently, it has been shown that under very low R:FR conditions phyA protein tends to accumulate [4, 8, 10].

SAS implementation is regulated by the interaction of active phyB with <u>P</u>HYTOCHROME INTERACTING <u>F</u>ACTORS (PIFs), a family of basic-helix-loop-helix (bHLH) transcription factors. When interacting with active phyB, PIFs are phosphorylated, which triggers the degradation of PIF1, PIF3, PIF4 and PIF5 (known as the PIF quartet, PIFQ) via the 26S proteasome. By contrast, PIF7 phosphorylation has little effect on its degradation but inhibits its DNA-binding activity [11]. Either case, under high R:FR (that mimics low plant density) PIF transcriptional activity is inhibited by the active form of phyB. Under low R:FR, active phyB is mostly eliminated, which results in PIF accumulation and/or promotion of their DNA-binding activity. This initiates a transcriptional cascade that leads to the expression of dozens of *<u>P</u>HYTOCHOME <u>R</u>APIDLY <u>R</u>EGULATED* (*PAR*) genes, several of which encode transcription factors from various families (e.g. bHLH, HD-Zip, BBX, …) having positive, negative or even complex roles in implementing the SAS hypocotyl [12–22]. Generally speaking, hypocotyls of seedlings deficient in negative and positive regulators elongate more and less, respectively, than those of wild-type seedlings when growing under simulated shade.

PIF7, together with a minor contribution of PIF4 and PIF5, has a major and positive role in promoting the shade-induced hypocotyl elongation. Consistently, *pif7* and the *pif4 pif5 pif7* (from now on *pif457*) showed an attenuated and almost null hypocotyl elongation in response to simulated shade [11, 23]. From the various *PAR* genes, induction of *YUCCAs* (*YUCs*) contributes to auxin production together with *SHADE AVOIDANCE 3* (*SAV3*, also known as *TRYPTOPHAN AMINOTRANSFERASE 1*/*WEAK ETHYLENE INSENSITIVE 8*, *TAA1*/*WEI8*) in the two-step indole-3-acetic acid (IAA) pathway from tryptophan (Trp). Indeed, the Trp aminotransferase encoded by *SAV3* catalyzes the conversion from Trp to Indole-3-pyruvic acid (IPyA), and the flavin monoxigenase encoded by *YUC* genes catalyzed the IPA oxidative decarboxylation to IAA (Figure 1) [24, 25]. Consistently, the single *sav3* and multiple mutants in *YUC* genes (*yuc2 yuc5 yuc8 yuc9* and *yuc3 yuc5 yuc7 yuc8 yuc9*) had attenuated shade-induced hypocotyl elongation [11, 12]. Together, these results highlight the importance of SAV3 and PIF-mediated *YUC* expression in this shade-mediated growth process.

Another *PAR* gene with a well-known negative role in the shade-induced hypocotyl elongation is *LONG HYPOCOTYL IN FR 1* (*HFR1*), that encodes a transcriptional cofactor of the bHLH family structurally related to PIFs but lacks the phyB- and DNA-binding ability [26, 27]. HFR1 inhibits the activity of all four PIFQ members (PIF1, PIF3, PIF4, PIF5) [28–30] and PIF7 [22, 31, 32] by heterodimerizing with them. Consistent with its negative role, *hfr1* hypocotyls are longer than wild-type ones under simulated shade [14, 18, 33]. A different case is *HYPOCOTYL ELONGATION 5* (*HY5*) known to encode a transcription factor of the basic domain-leucine zipper (bZIP) family. *HY5* expression is not as rapidly nor strongly induced in response to simulated shade (> 4 h of exposure to low R:FR) as that of the *PAR* genes described before (15-60 min). Shade-induced *HY5* expression is also phyA-dependent [18]. Hypocotyls of the *hy5* mutant seedlings elongate more than the wild-type ones under low R:FR [34–36], particularly to shade signals in response to inhibitory daily sunflecks, unfiltered light gaps that occur in the vegetation canopies, given in the afternoon [37]. Therefore, *HY5* acts as a negative SAS regulator. Recently it has been shown that shade avoidance is regulated by a three-layered gas-and-brake mechanism that involves a complex network of interacting bHLHs [22]. Nuclear-pore complex components, chloroplast derived signals and epigenetic components also prevent an excessive response to shade, providing additional levels of regulation of this response [35, 38, 39].

The mechanisms that work between SAS components to perceive and respond to simulated shade have been established in a few cases, as those between HFR1, PIFs and SAV3 [reviewed by [1, 2]]. In other cases, however, how some components work together is less clear. For instance, how HY5 exerts its negative effect in this pathway is unknown (Figure 1). The phyA accumulation in canopy shade prevents an excessive hypocotyl elongation by binding to and stabilizing the auxin signaling repressors AUX/IAA [10]. It is however unclear, whether phyA might also negatively regulates SAS modulating other well-established SAS regulators, such as PIFs (Figure 1).

To study these connections further, in this work we will explore the architecture of the SAS regulatory network. To do so, from the different regulators with a role in the shade-triggered hypocotyl elongation, we have focused in *PHYA*, *PHYB*, *HFR1* and *HY5* (negative regulators), *SAV3*, *PIF4*, *PIF5* and *PIF7* (positive regulators) because the available mutants of these genes show a clear alteration in the hypocotyl elongation in response to simulated shade (Figure 1). Using these mutants, we have taken different approaches to explore the relationships and connections between these SAS regulators: (1) pharmacological application of auxin-related inhibitors, to discover if the different negative regulators inhibit hypocotyl growth through regulating the same aspects of auxin activity, (2) genetic analyses, to establish if different SAS components work in the same or different regulatory branches or modules of the network; (3) temporal analyses, to learn when the different components analyzed act in controlling the shade-induced hypocotyl elongation; and (4) spatial analyses, to identify the cells targeted along the hypocotyl axis epidermis by each SAS regulator. Besides, we explore molecular connections between HY5 and PIFs, two antagonistic SAS components known to act both as transcription factors. Our findings indicated that these components are grouped in two main modules or branches that act at different times and impact the elongation of different cells along the hypocotyl axis. We also show that in these processes, the functional relationship between PIF457 and HY5 change with the time of shade exposure.

## RESULTS

### Mutant *phyA* and *hy5* seedlings respond to auxin inhibitors differently than *hfr1*

The cotyledons of *A. thaliana* and *Brassica rapa* seedlings perceive shade and trigger local IAA synthesis in the cotyledons themselves (Figure 1). The ability of auxin to travel over long distances facilitates the transport of IAA between the cotyledons and the hypocotyl, where cellular elongation occurs [40]. Consistently, treatments of wild-type seedlings with the auxin biosynthesis inhibitor L-kynurenine (L-kyn), that effectively and specifically targets Trp aminotransferases such as SAV3, (Figure 1) [41], or with the auxin transport inhibitor naphthylphthalamic acid (NPA) abolish the hypocotyl elongation response in wild-type seedlings [16, 25, 42, 43]. In our W+FR conditions, the extra elongation of *hfr1* compared to wild-type hypocotyls was completely abolished by the highest doses of L-kyn applied (Figure 2A). Auxin quantification indicated that IAA levels in Col-0 and *hfr1* seedlings increased to similar values after 1 h of W+FR treatment (Figure 2B), which suggests that HFR1 does not have a strong and measurable impact on the IAA levels, at least at the time of shade treatment analyzed. By contrast, the extra hypocotyl elongation of *phyA* and *hy5* seedlings was much less affected by L-kyn than the wild type, even at the highest concentration tested (Figure 2A). In addition, IAA levels in *phyA* were significantly reduced even before the shade treatment; they were also significantly attenuated after 1 h of W+FR in both *hy5* and *phyA* (Figure 2B). These results suggested that shade-induced elongation in these two mutant lines was not fully dependent on auxin levels. As IAA levels seem to be under negative feedback control, as in several auxin-signaling mutant [44, 45], these results are consistent with *phyA* and *hy5* having an altered auxin responsiveness [10, 46].

**Figure 2.**
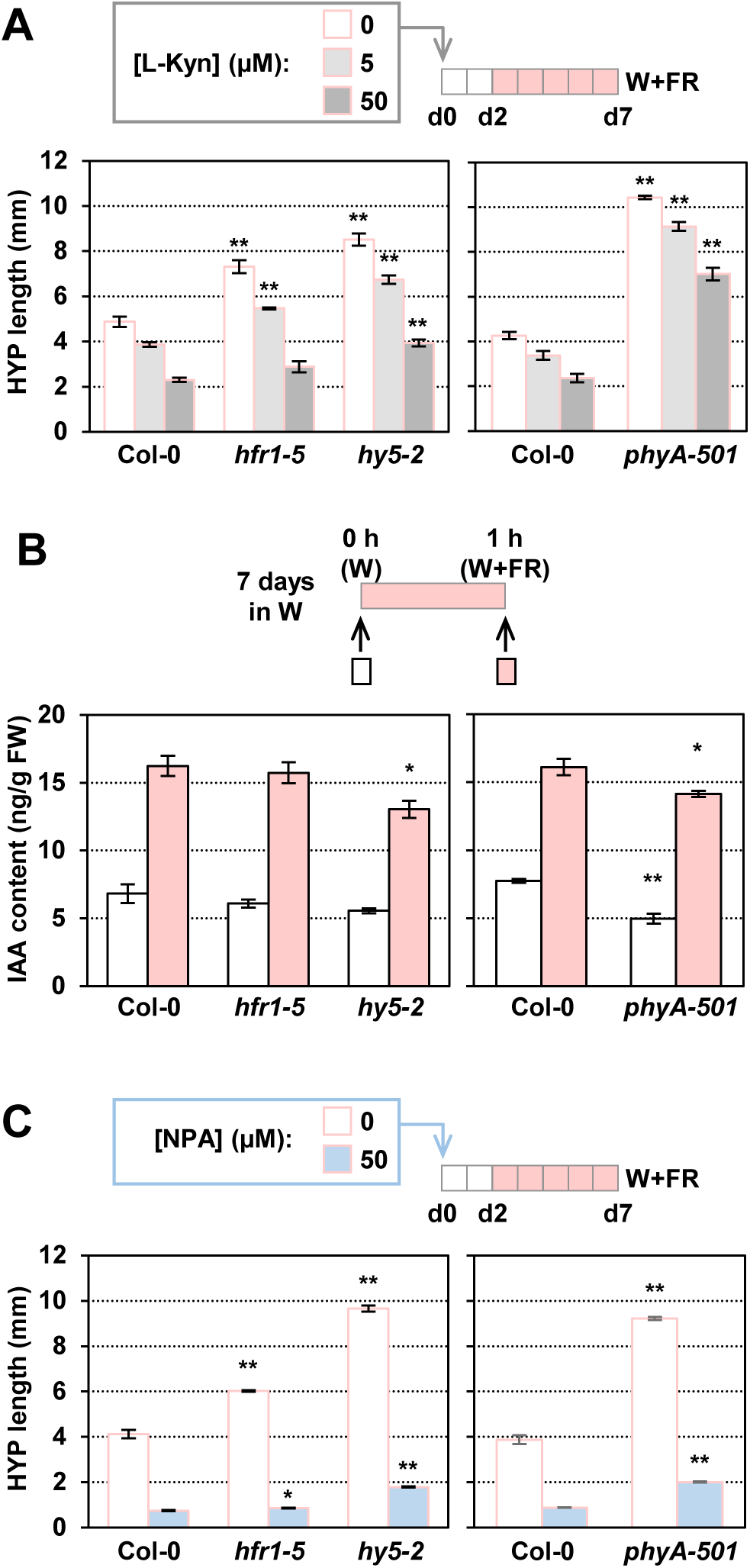
Contribution of auxin synthesis and polar transport in the shade-induced hypocotyl response of *phyA*, *hy5* and *hfr1* seedlings. **(A)** Effect of increasing concentrations of L-kynurenin on the shade-induced hypocotyl length of Col-0, *hfr1-5*, *hy5-2* and *phyA-501* seedlings. Seedlings were grown in W for 2 days and then transferred to W+FR for 5 more days. **(B)** IAA content in Col-0, *hfr1-5*, *hy5-2* and *phyA-501* seedlings grown in W for 7 d and then treated for 1 h with W+FR (R:FR = 0.02). Whole seedlings were collected to measure IAA levels. Data are presented as means and SE of four biological replicates. FW, fresh weight. **(C)** Effect of NPA on the shade-induced hypocotyl length of Col-0, *hfr1-5*, *hy5-2* and *phyA-501* seedlings. Seedlings were grown in W for 2 days and then transferred to W+FR for 5 more days. In **A** and **C**, inhibitors were applied in the media. Asterisks indicate significant differences (Student’s *t* test) relative to the wild-type growing under the same light treatment (ns, not significant, * P-value < 0.05, ** P-value < 0.01).

The extra elongation of *hfr1* mutant seedlings is almost abolished by the application of the auxin transport inhibitor NPA (Figure 2C), which indicates that the action of PIFs-HFR1 in modulating the shade-induced hypocotyl elongation is mostly dependent on auxin produced somewhere else and transported to the hypocotyl [as proposed [42]]. By contrast, *phyA* and *hy5* showed a clear resistance to the inhibitory effect of NPA (Figure 2B), which suggested that these two factors share similar mechanisms to repress shade-induced elongation and, in contrast with HFR1, are less dependent on auxin transport to promote elongation.

### The SAS regulatory network is organized in at least two genetically differentiated modules or branches

We first prepared a series of genetic crosses focusing on a few mutants in negative (phyA, phyB, HY5, HFR1) or positive (SAV3, PIF4, PIF5 and PIF7) SAS components (Figure 1). These mutants result in strong shade-related hypocotyl phenotypes. From these components, only PIF4, PIF5 and PIF7 (PIF457) are known to show some redundancy in controlling the shade-induced hypocotyl elongation [11, 42]. The hypocotyl length of the single and double phyA and phyB mutant seedlings in response to simulated shade was first analyzed (Figure 3). In white light (W) (that simulates sunlight of high R:FR), the length of Col-0 and *phyA* hypocotyls was similar, whereas that of *phyB* hypocotyls was longer and those of *phyA phyB* double mutant seedlings were the longest, as expected. In W+FR (very low R:FR), *phyA* hypocotyls were longer and *phyB* hypocotyls shorter, respectively, than the wild type (Figure 3A, B). The antagonistic activity of phyA and phyB under simulated shade indicate that the W+FR conditions employed in these experiments mimic canopy shade, in contrast with those proximity shade conditions in which phyA action is negligible [4]. Importantly, the *phyA phyB* hypocotyl length in W+FR was even longer than in W (Figure 3B), in agreement with other phytochromes regulating the shade-induced hypocotyl elongation [47]. To visualize better the effect of simulated shade in controlling hypocotyl elongation, particularly useful when comparing genotypes with different hypocotyl length under W (e.g., Col-0 vs. *phyB*), the difference in hypocotyl length in W+FR and W (HYP_W+FR_ - HYP_W_) was calculated (Figure 3C). This representation showed that *phyA phyB* double mutant hypocotyls had an intermediate shade-induced elongation response that those of *phyA* and *phyB* single mutants (Figure 3C), suggesting that the effect of the two phytochromes is additive, likely acting independently of one another in controlling the shade-induced hypocotyl length. In the following set of experiment, the HYP_W+FR_ - HYP_W_ will be shown when comparing the different mutants (the hypocotyl length data will be included as supplemental information).

**Figure 3.**
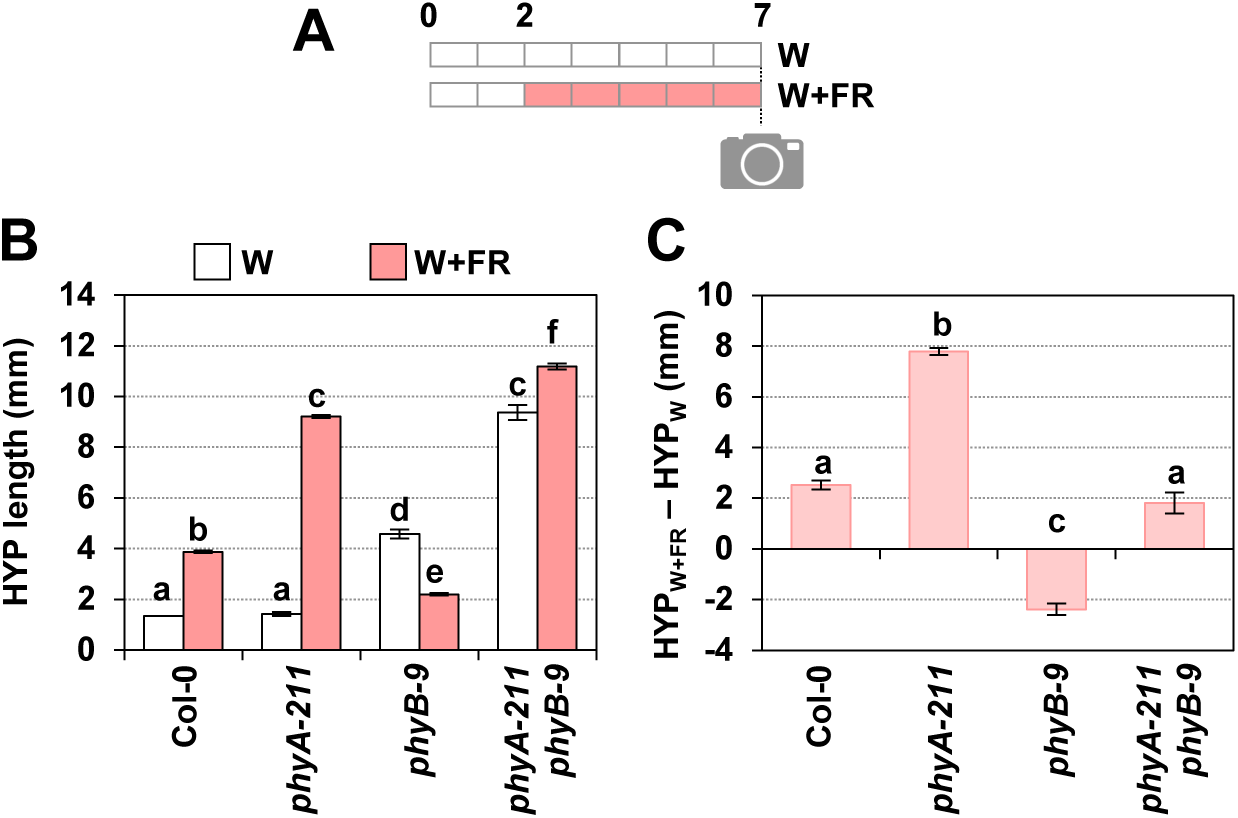
Genetic interaction of phyA and phyB in the regulation of the shade-induced hypocotyl elongation. **(A)** Cartoon showing the design of the experiment. Seedlings were germinated and grown in white light (W, R:FR > 1.5) for 2 days and then they either maintained in W or transferred to simulated shade (W+FR, R:FR, 0.02) for 5 more days. On day 7, hypocotyl length was measured. **(B)** Hypocotyl length of Col-0, *phyA-211*, *phyB-9* and *phyA-211 phyB-9* double mutant seedlings after growing in W or W+FR. Values are means and SE of three independent samples. **(C)** Elongation response of seedlings of the lines shown in **B**. The means of HYP_W_ and HYP_W+FR_ (data shown in **B**) were used to calculate HYP_W+FR_-HYP_W_. SE were propagated accordingly. Different letters denote significant differences (one-way ANOVA with the Tukey test, P-value < 0.05) among means.

We next produced double mutants deficient in other negative regulators (*phyA hfr1*, *hy5 hfr1* and *phyA hy5*) and analyzed their shade-induced hypocotyl elongation response (no *phyB* mutant was included in these crosses as its phenotype was observed less clearly in W+FR) (Figure 4, Supplementary Figure S1). Seedlings of *phyA hfr1* and *hy5 hfr1* double mutant elongated more than the single mutants (Figure 4A-B), suggesting that they worked additively, in agreement with previous information [18]. By contrast, *phyA hy5* seedlings elongated as much as the *phyA* single mutant (Figure 4C), indicating that phyA was epistatic over HY5. Together, our genetic analyses suggested that (1) phyA and HY5 act in the same branch of the SAS regulatory network; and (2) HFR1 acts independently of phyA and HY5 in controlling this response. Therefore, it seems that these components are likely organized in two separate branches or modules, one of which involves the action of phyA and HY5 but not HFR1.

**Figure 4.**
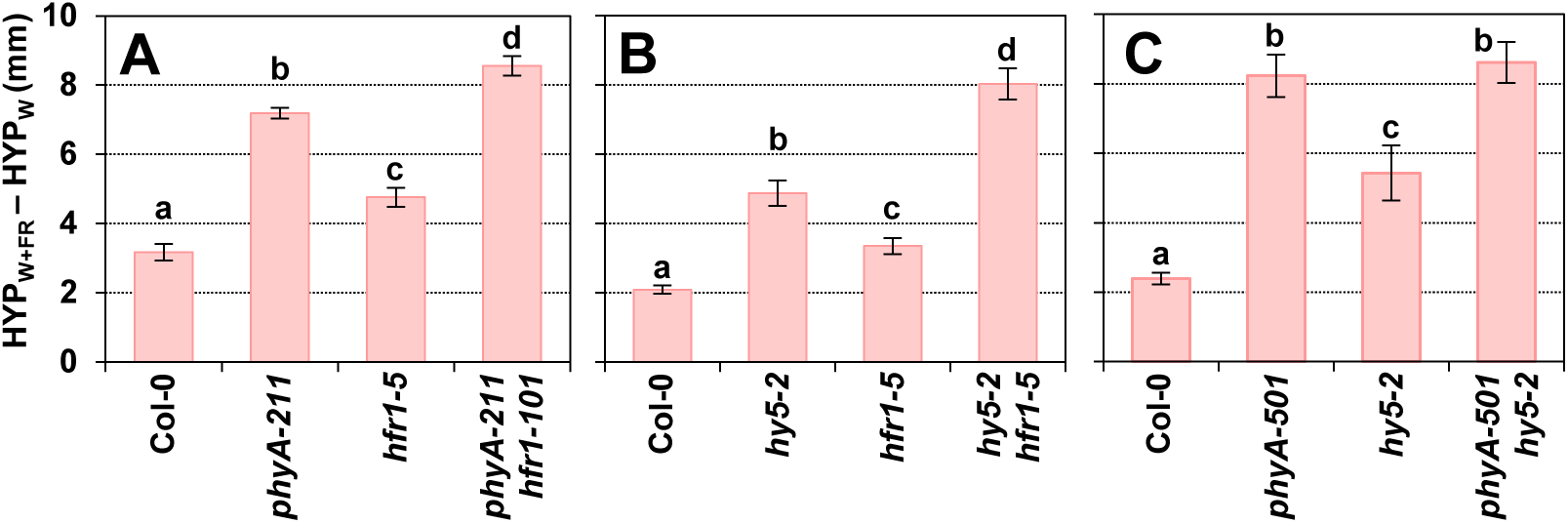
Genetic interaction of SAS negative regulators in the regulation of the shade-induced hypocotyl elongation. Difference of hypocotyl length in W+FR (HYP_W+FR_) and W (HYP_W_) of Col-0, **(A)** *phyA-211*, *hfr1-5, phyA-211 hfr1-101*, **(B)** *hy5-2*, *hfr1-5*, *hy5-2 hfr1-1*, **(C)** *phyA-501, hy5-2* and *phyA-501 hy5-2.* Seedlings were germinated and grown as indicated in Figure 3A. Values are means and SE of three independent samples. Different letters denote significant differences (one-way ANOVA with the Tukey test, P-value < 0.05) among means.

We also generated multiple mutants between positive and negative SAS regulators (Figure 5; Supplementary Figure S2). As before, *phyB* mutants were excluded from these crosses. Phenotypic analyses showed that *hfr1 pif7* length was intermediate between the single mutants (Figure 5A; Supplementary Figure S2A) consistent with HFR1 interacting with and inhibiting PIF7 activity [7, 22, 31]. Because HFR1 is known to heterodimerize with and inhibit the transcriptional activity of several PIFs [11, 26, 28, 29, 48], we reasoned that the intermediate *hfr1 pif7* elongation was likely caused by the presence of the SAS positive regulators PIF4 and PIF5. Consistently, the elongation response to shade of hypocotyls of an *hfr1 pif457* quadruple mutant line was reduced compared to *hfr1 pif7*, although not completely abolished and still longer than *pif457* seedlings (Figure 5B; Supplementary Figure S2B), reflecting the minor contribution of additional factors. Mutant *hfr1* plants were also crossed with *sav3*, known to have a role in the shade-induced auxin biosynthesis [49]. Importantly, phenotypic analyses showed that *hfr1 sav3* (Figure 5C; Supplementary Figure S2C) was almost as short as *hfr1 pif457* (although significantly longer than *sav3* seedlings), in agreement with the described role of PIF457 in promoting the SAV3-dependent IAA biosynthesis [11, 27].

**Figure 5.**
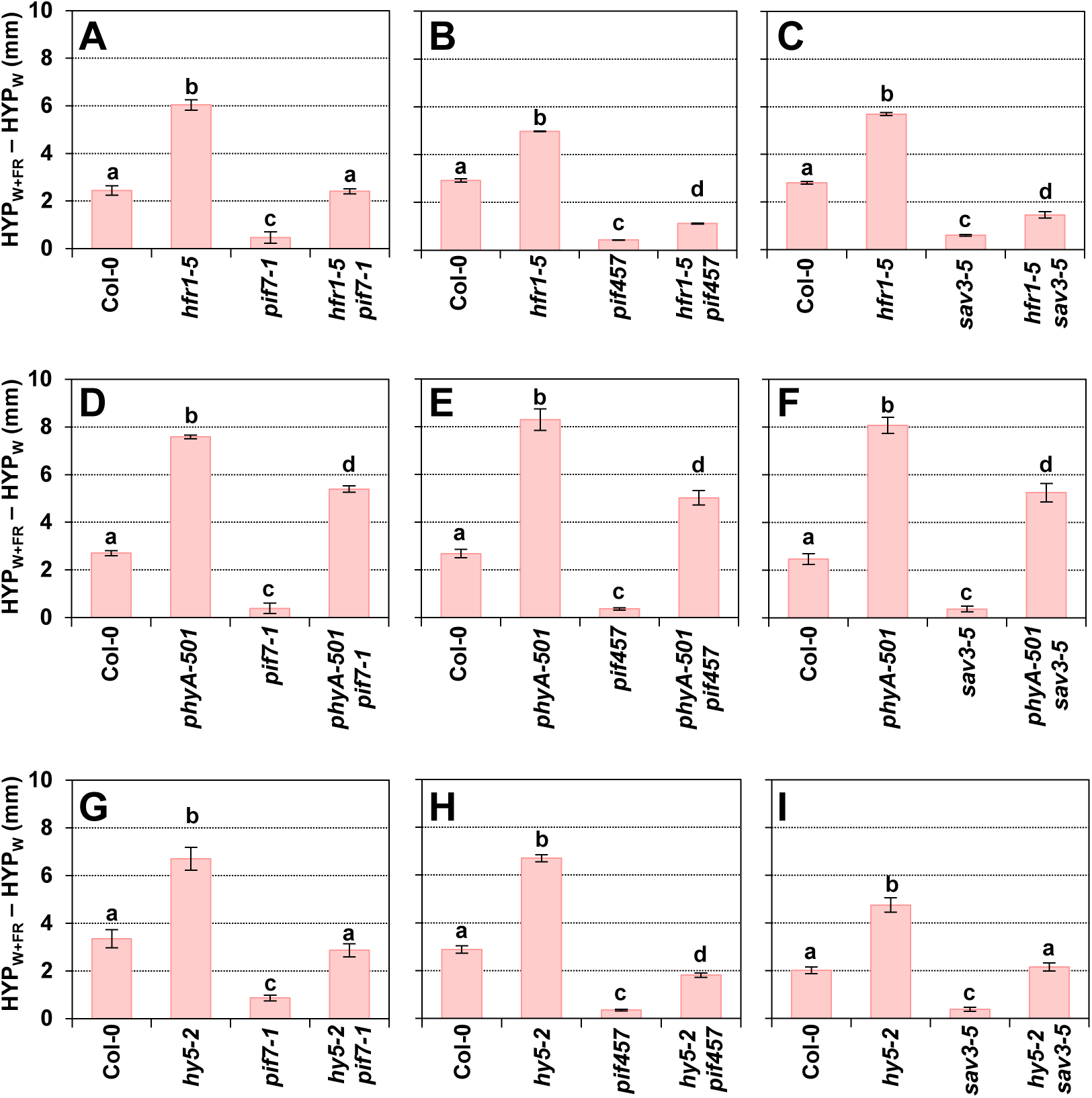
Genetic interaction of pairs of SAS negative and positive regulators in the regulation of the shade-induced hypocotyl elongation. Difference of hypocotyl length in W+FR (HYP_W+FR_) and W (HYP_W_) of Col-0 **(A)** *hfr1-5*, *pif7-1*, *hfr1-5 pif7-1*, **(B)** *hfr1-5*, *pif457*, *hfr1-5 pif457*, **(C)** *hfr1-5*, *sav3-5*, *hfr1-5 sav3-5*, **(D)** *phyA-501*, *pif7-1*, *phyA-501 pif7-1*, **(E)** *phyA-501*, *pif457*, *phyA-501 pif457*, **(F)** *phyA-501*, *sav3-5*, *phyA-501 sav3-5*, **(G)** *hy5-2*, *pif7-1*, *hy5-2 pif7-1*, **(H)** *hy5-2*, *pif457*, *hy5-2 pif457*, and **(I)** *hy5-2*, *sav3-5*, *hy5-2 sav3-5.* Seedlings were germinated and grown as indicated in Figure 3A. Raw data are shown in Supplemental Figure S2. Values are means and SE of three independent samples. Different letters denote significant differences (one-way ANOVA with the Tukey test, P-value < 0.05) among means.

Mutant *phyA* plants were next crossed with *pif7* or *pif457*. In contrast with the previous crosses, *phyA pif7* and *phyA pif457* shade-induced hypocotyl elongation was similar between them and closer in length to that of *phyA* than to *pif7* or *pif457* (Figures 5D-E). A similar elongation response was observed in the *phyA sav3* seedlings (Figures 5F). Together, these results suggested that the elongation repression imposed by phyA under simulated shade is mostly independent on PIF457 or the rapid shade-induced and SAV3-dependent auxin biosynthesis (Figures 5D-F; Supplementary Figure S2D-F).

Finally, *hy5* plants were crossed with *pif7* and *pif457*. Hypocotyls of *hy5 pif7* and *hy5 pif457* showed an intermediate elongation that the parental *pif7*, *pif457* and *hy5* lines (Figures 5G-H). A similar intermediate pattern of shade-induced elongation was observed for the *hy5 sav3* seedlings (Figure 5I). Together, these results suggest an additive activity for these regulators in the control of this shade-induced elongation response (Figure 5G-I; Supplementary Figure S2G-I). These conclusions are consistent with a network architecture in which PHYA and HY5 act independently of PIF457, HFR1 and SAV3 to promote the shade-induced hypocotyl elongation.

### SAS regulatory components act in different moments during the shade-induced hypocotyl elongation

To learn if the various SAS components act at the same time or in different moments after the exposure of young seedlings to W+FR, growth rates were first determined in wild-type (Col-0) seedlings grown under W and W+FR. To do so, hypocotyl length was measured daily from day 2 to day 7 in different groups of seedlings, and the variations in the daily growth rate were estimated for each genotype and light treatment (Figure 6A). Under W, Col-0 growth rate remained low but constant along the period investigated (Figure 6B, D), whereas under W+FR it went up from day 5 onwards (Figure 6C, E). As an additional control, growth rate of the *phyB* mutant hypocotyls was also estimated. Importantly, under W, *phyB* growth rate increased with the age of the seedlings, elongating more at the end of the period analyzed (from day 5 onward) (Figure 6B). Under W+FR, *phyB* growth rate mimicked that of Col-0 but it was attenuated, with higher growth rate at the end of the period (Figure 6C), consistent with its reduced elongation compared to Col-0 (Figure 3) [4, 8]. Under W, growth rate of *phyA*, *hy5* and *hfr1* hypocotyls was quite constant along time and similar to those of Col-0, except on day 2, when phyA hypocotyls grew slightly faster than Col-0 (Figure 6B, D). Under W+FR, *phyA* growth rate was much higher than that of Col-0 hypocotyls on days 2-4 (peak on day 3) but progressively dropped to values closer to those of Col-0 on day 6 (Figure 6C, E). Similarly, *hy5* grew faster at the beginning of the development although its growth rate peaked on day 4 (Figures 6E). By contrast, *hfr1* also had a peak of growth on day 4 but elongated faster in the second half of the period of time analyzed (Figures 6E).

**Figure 6.**
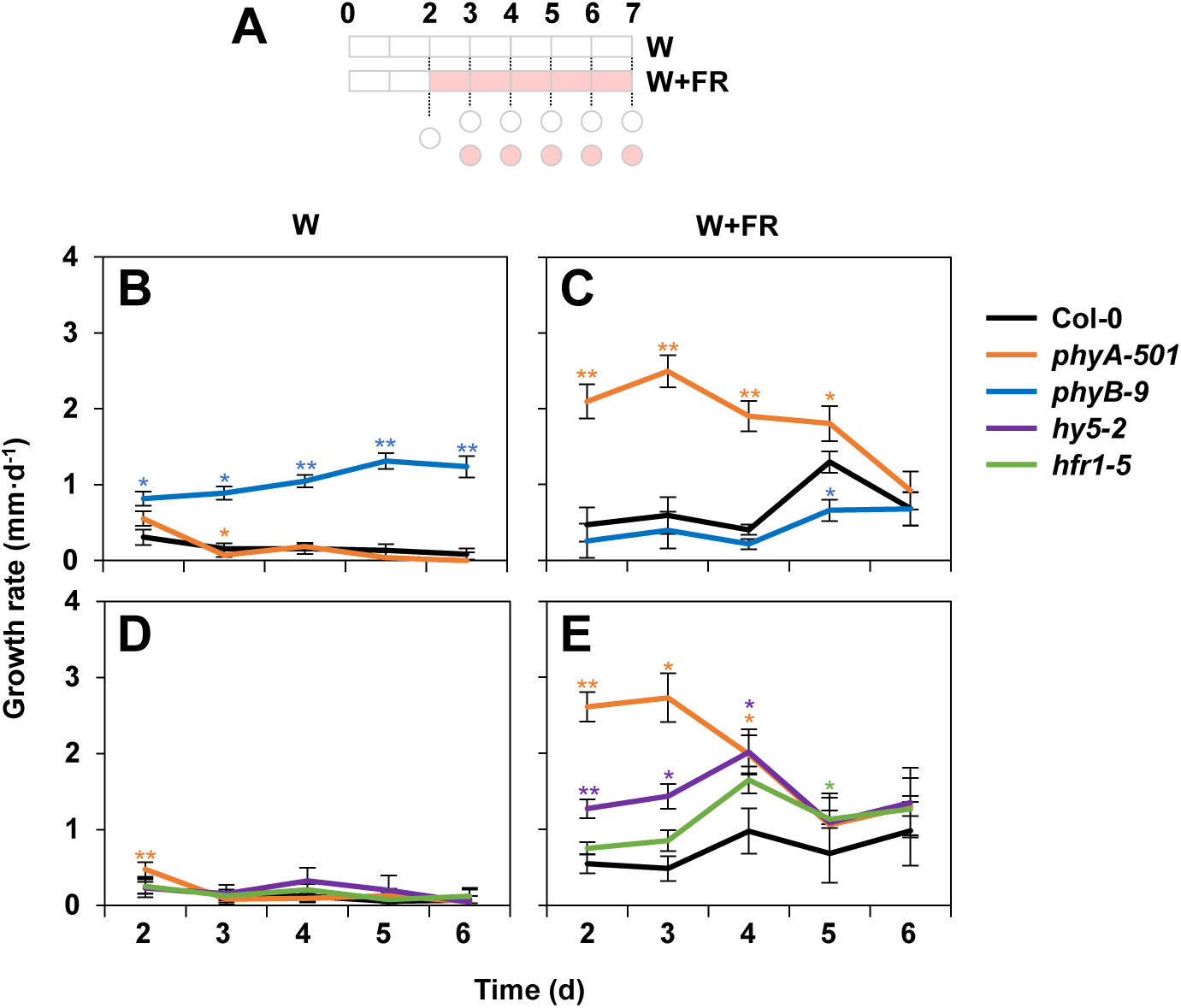
Effect of phyA, phyB, HFR1 and HY5 mutations on the growth rate of the hypocotyl under W and W+FR. **(A)** Cartoon of the experiment design. Seeds were germinated and grown for 2 days under W and then either kept under W or transferred to W+FR (R:FR = 0.02) for 5 more days. Circles indicate the days on which hypocotyls were measured to estimate the daily growth rate (difference of hypocotyl length per day). Growth rate of Col-0, *phyA-501* and *phyB-9* hypocotyls grown in W **(B)** or in W+FR **(C)**. Growth rate of Col-0, *phyA-501*, *hfr1-5* and *hy5-2* hypocotyls grown in W **(D)** or in W+FR **(E)**. Values have been estimated after daily mean lengths and SE of three independent biological replicates per day, genotype and light treatment.

To visualize the repressor activity of the different regulators, the growth rate of the wild-type was subtracted to that of each mutant grown in those conditions where the phenotype is more obvious: W for *phyB*, and W+FR for *hfr1*, *hy5* and *phyA*. This representation confirmed our previous conclusions (Supplementary Figure S3). In summary, although the temporal activity of the regulators overlapped, phyA and HY5 repressed hypocotyl elongation more strongly at the beginning of seedling development (from days 2 to 4) and differed from HFR1 (and phyB), whose activity was more important at the second half of the period analyzed (from days 5 to 7). These results provided a temporal framework that separates the action of the participating components.

### SAS regulatory components target overlapping but different regions along the hypocotyl axis

In *A. thaliana*, hypocotyl elongation is a result of cell elongation (not cell division). Among the different tissues of this organ, the epidermis is of particular importance in mediating auxin-induced growth in the hypocotyl [50]. The pattern of epidermal cell length in W-grown hypocotyls was shown to take place in all cells over the entire growth period (from 1 to 9 days after germination), although the area of fastest growth moves acropetally, from the base (cells 2-4) on days 1-2 to the middle (cells 10-12) of the hypocotyl on days 7-9 [51]. Growth in dark-grown seedlings also initiates in the hypocotyl basal cells but, in this case, cells that elongated fastest move up much more rapidly and only a few cells upwards: from cell 1 at 36-48 h to cells 3-4 at 72 h from germination [51]. As there is not much information about how *A. thaliana* hypocotyls elongate in response to simulated shade at the cell level, we first established the pattern of cell elongation in wild-type (Col-0) hypocotyls grown under W and W+FR. Using confocal microscopy, the length of several files of epidermal cells along the hypocotyl longitudinal axis per treatment were measured (Supplementary Figure S4). Values were averaged for each of the about 20 cells that constitute a cell file (from bottom to top) along the hypocotyl longitudinal axis. Cell length in W-grown hypocotyls was similar along the hypocotyl (Supplementary Figure S5). By contrast, W+FR treatment enhanced the elongation of all cells compared to W treatment, although the pattern of cell elongation was not uniformly distributed, with cells located in the lower half of the hypocotyl elongating the most (Supplementary Figure S5). A similar conclusion was reached when representing the difference in length between cells grown in W+FR and W in each position, with cells 7-8 being the ones that grew the most, becoming about 170-250 µm longer than cells in the same position of W-grown hypocotyls (Figure 7, Col-0 panels). These results indicate that the shade-induced hypocotyl elongation of wild-type seedlings resulted from a bell-shaped non-symmetrical (skewed) elongation pattern of epidermal cells peaking around cells 7-8 from the base.

**Figure 7.**
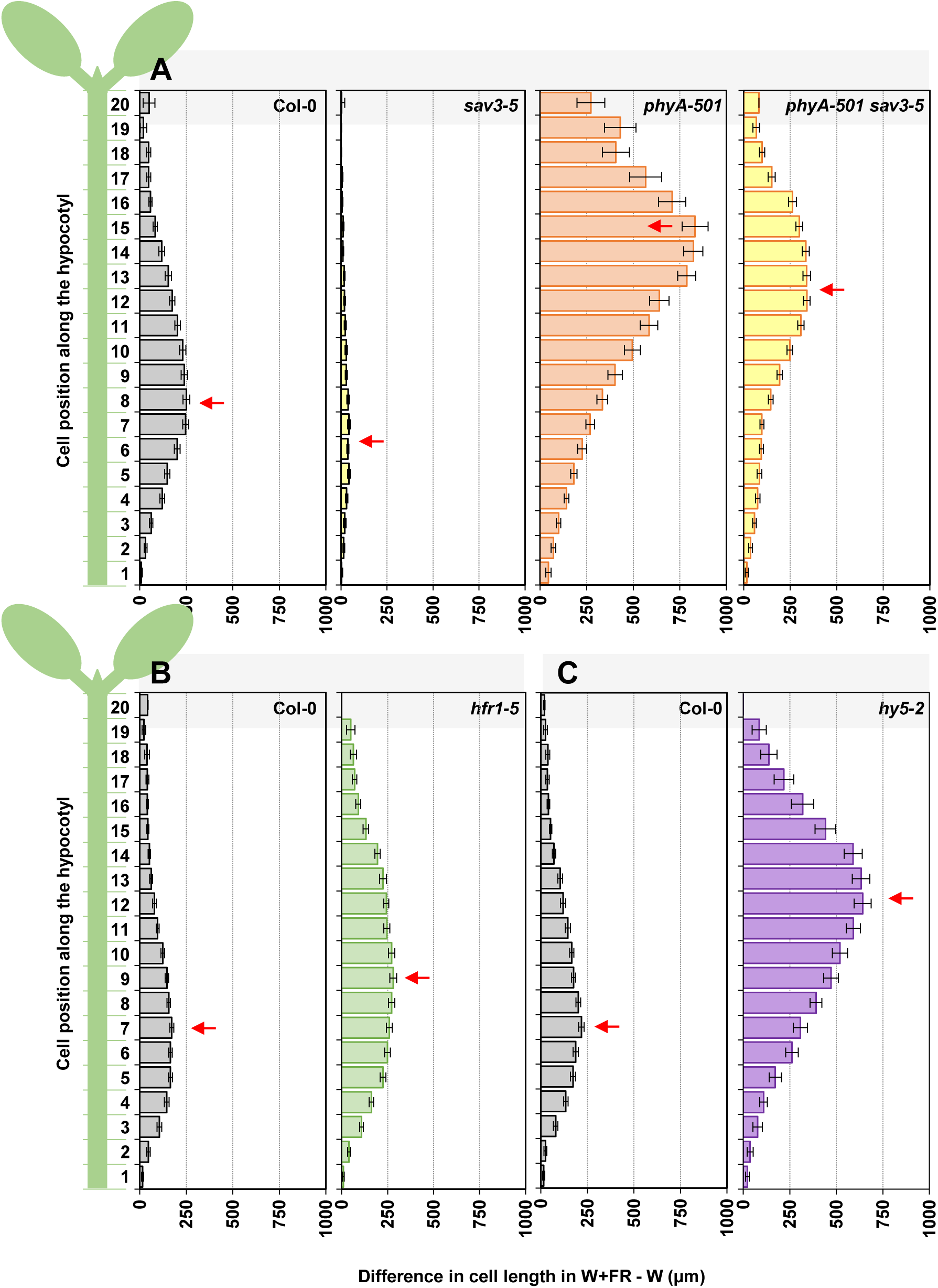
Distribution of the epidermal cell length from the base to the top induced by simulated shade in hypocotyls of wild type seedlings or deficient in SAS regulators. A schematic representation of the 20 cells that comprise an epidermis cell file along the longitudinal axis of an *A. thaliana* hypocotyl is shown at the left of graphs. Difference in length in W+FR and W for each of the 20 epidermal cells (from bottom to top) that conform a file in hypocotyls of Col-0, (**A)** *sav3-5*, *phyA-501*, *phyA-501 sav3-5*, **(B)** *hfr1-5* and **(C)** *hy5-2*. Seedlings were germinated and grown as indicated in Figure 3A. Red arrows point to the cell that presents a higher difference in length. Cell lengths in W and W+FR are shown in Supplementary Figure S5. Values have been estimated after mean lengths and SE of at least 15 cells of 2 cell files per hypocotyl from 7 different hypocotyls per genotype and growth condition.

Independently on the primary site of action of the studied SAS regulators (e.g., cotyledons or hypocotyls), their activities converge on the elongation of hypocotyls. To establish whether the convergence affected the same or different hypocotyl cells, we next analyzed the shade-induced cell length in hypocotyls of seedlings deficient in specific SAS regulators (Supplementary Figure S5, Figure 7). The hypo-responsive *sav3* hypocotyls showed a similar pattern of cell elongation as wild type (i.e., bell-shaped non-symmetrical) but strongly attenuated and slightly shifted to lower cells (elongation peak in cells 5-7 that elongated ∼40 µm more than the same cells in W-grown hypocotyls) (Figure 7A). In the shade-hyper-responsive *hfr1* seedlings, the peak of cell length was shifted closer to the middle part of the hypocotyl, with cell 9 showing the maximum of elongation (∼280 µm longer than cell 9 in W-grown hypocotyls) (Figure 7B). In the case of *hy5* and *phyA*, also hyper-responsive to shade, cell length was strongly enhanced and the elongation peak moved to the upper half of the hypocotyl (cell 12 in *hy5* that elongated ∼640 µm more than the same cell in W-grown hypocotyls; cell 15 in *phyA* that elongated ∼830 µm more than the same cell in W-grown hypocotyls) (Figure 7A, C).

As the peak cell number correlated with the difference in hypocotyl length in W+FR and W (HYP_W+FR_ - HYP_W_) (Supplementary Figure S6), we wondered if the redistribution of cell growth was a consequence of the enhanced hypocotyl shade-induced elongation shown by these genotypes. To check this possibility, we analyzed the cell length in *phyA sav3* hypocotyls, whose shade-induced hypocotyl elongation was similar to that of *hfr1* and shorter than *hy5* hypocotyls (Figure 6). In *phyA sav3* seedlings, the peak of cell elongation was similar to that of *hy5* and *phyA* (elongation peak in cell 13) and longer (∼340 µm longer than cell 13 in W-grown hypocotyls) than the longest cell in *hfr1* (cell 9, ∼280 µm longer than same cell in W-grown hypocotyls), reinforcing that phyA represses the elongation of a group of cells located in the upper half of the hypocotyl (Figure 7A). Consistently, in this case peak cell number did not correlate with the HYP_W+FR_ - HYP_W_ (Supplementary Figure S6).

Altogether, these analyses indicate that (1) the hypocotyl cells more responsive to simulated shade are located in the lower half of the wild-type hypocotyls (centered in cells 7-8), (2) deficiency in SAS negative regulators keeps the pattern of cell elongation but affects the peak cell number, and, (3) although the target cells of the various SAS negative regulators overlap, the peak cell number due to loss of *HY5* and *PHYA* function is strongly shifted towards the upper half of the hypocotyl. These results are consistent with phyA and HY5 activities repressing the cells of the upper part of the hypocotyl whereas HFR1 more clearly repressed the elongation of cells located in the lower half of the hypocotyl, providing a spatial framework that separates the action of the participating components.

### PIF457 and HY5 modulate the expression of shared shade-regulated genes

Despite the temporal and spatial differences observed between phyA/HY5 and PIFs/HFR1/SAV3 in controlling the hypocotyl cell elongation, their activities overlap and eventually converge. Hence, we aimed to further investigate possible convergence points between these two groups of regulators. Evidence in other photomorphogenic or temperature-regulated responses showing that HY5 directly interacts with PIF1/PIF3 proteins [52] and HY5 and PIF activities converge at a shared *cis* regulatory element [32, 53, 54] led us to explore shade-induced changes of PIF457, and HY5 in the expression of shared targets genes.

Amongst the >3000 genes identified as potentially putative HY5 binding targets [55], we focused on *1-AMINO-CYCLOPROPANE-1-CARBOXYLATE SYNTHASE 8* (*ACS8*) and *PAR1*. Both were also described as direct PIFs targets [10, 56, 57], therefore appearing as potential candidates for the convergence of HY5 and PIF457 transcriptional activities. Next, we compared the relative expression of these two genes in 7 days-old seedlings of Col-0, *hy5*, *pif457* and *hy5 pif457* exposed with W+FR for 0, 1, 4, 8 and 24 h (Figure 8A). The expression of both genes was significantly promoted in Col-0 after 1-8 h of W+FR treatment (compared to the beginning of the treatment) and decreased after 24 h of the shade treatment, reaching comparable levels to those found at 0 h. In *hy5*, *ACS8* and *PAR1* expression was also induced after 1-8 h and only *PAR1* expression levels were significantly higher than in Col-0. By contrast, in *pif457* and *hy5 pif457* the expression of *ACS8* and *PAR1* remained virtually unaffected by the W+FR treatment (Figure 8B). These results suggest that (1) PIF457 activates whereas (2) HY5 repress *PAR1* (and likely *ACS8*) expression. Importantly, (3) HY5 activity depends on PIF457 transcriptional activation.

**Figure 8.**
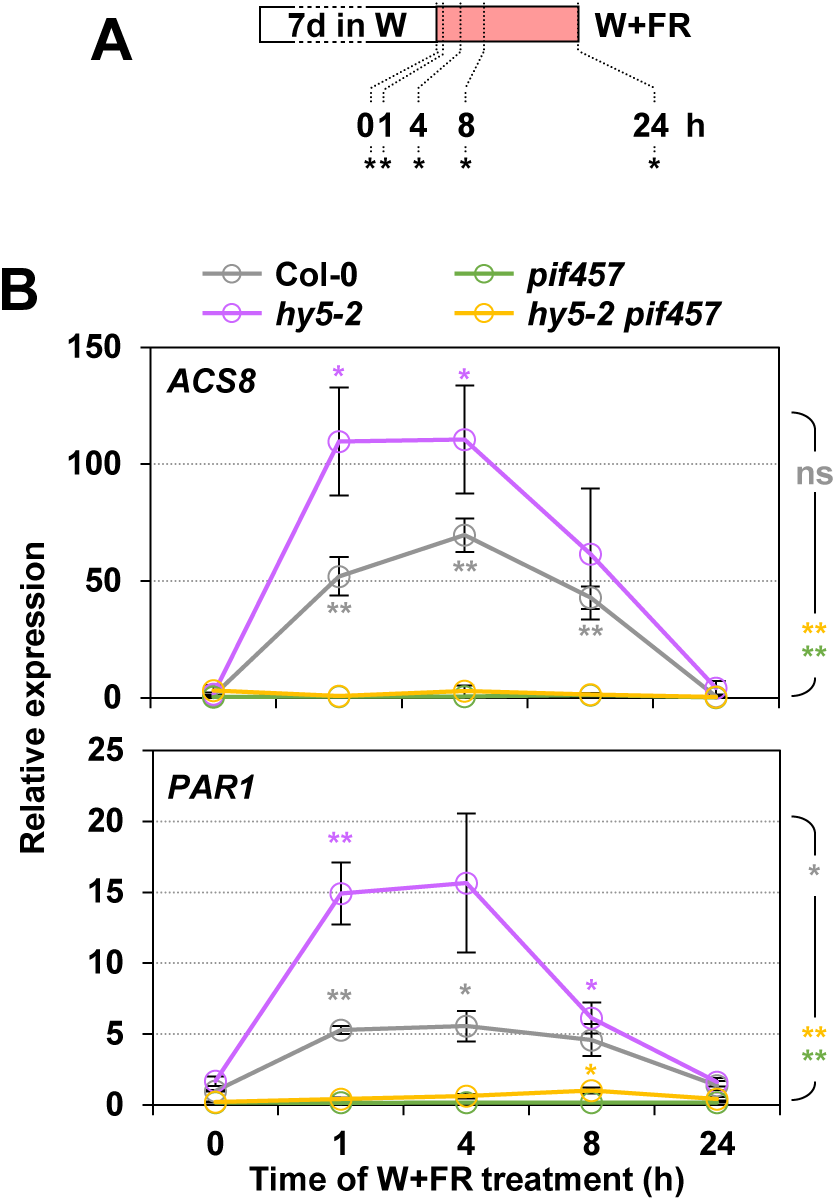
Effect of *hy5* and *pif457* mutations on the shade-regulation of the expression of common target genes. **(A)** Cartoon of the experiment design. Seeds of the indicated genotypes were germinated and grown for 7 days under W and then transferred to W+FR (R:FR = 0.02) for the indicated time periods before harvesting samples (indicated with asterisks). **(B)** Relative expression of *ACS8* (top panel) and *PAR1* (bottom panel) in Col-0, *hy5-2*, *pif457* and *hy5-2 pif457* at the indicated times of simulated shade treatment. Values are means and SE of three independent biological replicates relative to the Col-0 genotype at 0 h. Asterisks around the symbols indicate significant differences (Student’s *t* test) relative to the same genotype at 0 h. Asterisk at the right indicate significant differences between the different mutants and the wild-type in response to simulated shade (two-way ANOVA); ns, not significant, * P-value < 0.05, ** P-value < 0.01.

Next, we carried out RNA sequencing (RNA-seq) of the time points 0, 1 and 8 h after shade exposure of the four genotypes (Col-0, *hy5*, *pif457* and *hy5 pif457*) to expand our understanding of the role and interaction of HY5 and PIF457 activities in the transcriptomic dynamics early (0 h vs. 1 h) and late (0 h vs. 8 h) after shade treatment (Figure 9A). We identified differentially expressed genes (DEGs) up-(fold change ≥ 1.5, P < 0.05) and downregulated (fold change ≤ -1.5, P < 0.05) after 1 and 8 h of shade treatment compared with 0 h for each genotype analyzed (Supplementary Tables S1-S4).

**Figure 9.**
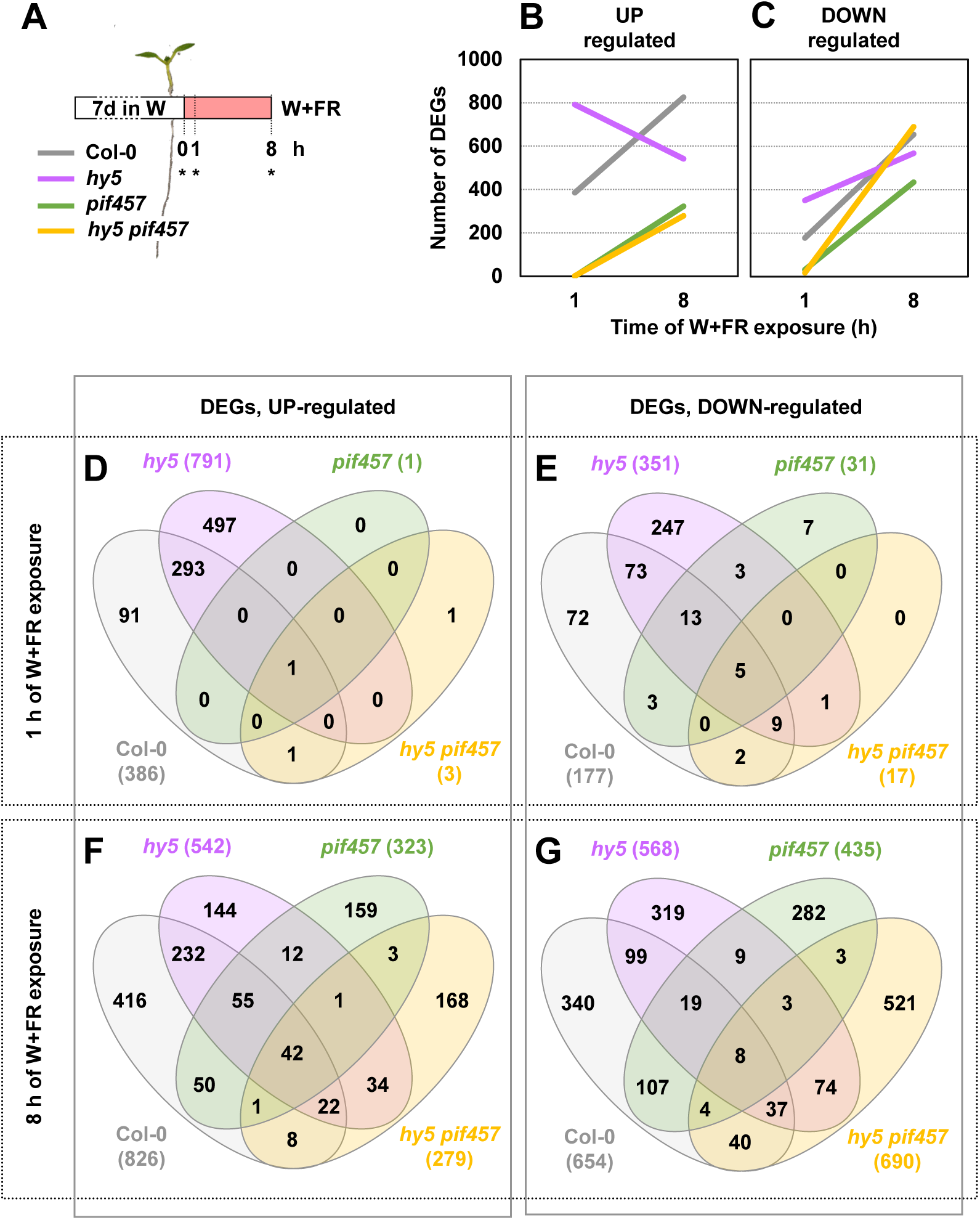
Effect of *hy5* and *pif457* mutations on the shade-regulated transcriptome. **(A)** RNA-seq was performed with RNA extracted from Col-0, *hy5-2*, *pif457* and *hy5-2 pif457* seedlings at the indicated times of simulated shade treatment. Seedlings were grown as indicated in Figure 8A. Asterisks indicate the harvesting moment for RNA extraction. Three independent biological replicates were used for each genotype and treatment. Evolution of the number of up- **(B)** and down-regulated **(C)** DEGs in response to 1 and 8 h of W+FR in Col-0, *hy5-2*, *pif457* and *hy5-2 pif457* seedlings grown as indicated in section **A**. Venn diagrams showing the overlap of up- **(D, F)** and down-regulated **(E, G)** DEGs after 1 **(D, E)** and 8 h **(F, G)** of W+FR treatment between Col-0, *hy5-2*, *pif457* and *hy5-2 pif457* seedlings.

After 1 h of W+FR, 386 and 791 DEGs were induced in wild-type and *hy5* seedlings, respectively. Only 1 and 3 genes were induced after 1 h of W+FR in *pif457* and *hy5 pif457* seedlings, respectively (Figure 9B, D). Regarding the down-regulated genes, 177 DEGs were repressed in wild-type seedlings. As before, an increased number of genes were repressed after 1 h of W+FR in *hy5* (351) and a reduced number in *pif457* (31) and *hy5 pif457* (17) seedlings (Figure 9C, E). From the total number of these early shade-modulated DEGs, 294 upregulated genes were shared between *hy5* (out of 791 genes, 37.2 %) and Col-0 (out of 386 genes, 76.2 %) (Figure 9D) and 100 downregulated genes where shared between *hy5* (out of 351, 28.5 %) and Col-0 (out of 674, 14.8 %) (Figure 9E). As 748 DEGs appeared only in *hy5* but not in Col-0 (497 upregulated and 251 downregulated), we concluded that HY5 has an important dual role as both activating and repressing the rapid changes in gene expression modulated by shade. Importantly, the vast majority of these genes did not appear as DEGs in *pif457* (1 upregulated or 31 downregulated) and *hy5 pif457* (3 upregulated or 17 downregulated) (Figure 9D, E), indicating that PIF457 are basically required for all the changes in gene expression (both activation and repression and either dependent or independent of HY5) that take place after 1 h of simulated shade exposure.

After 8 h of W+FR, 826 and 542 genes were induced in Col-0 and *hy5* seedlings, respectively. After this time of W+FR exposure, a reduced but substantial number of DEGs were activated in *pif457* (323) and *hy5 pif457* (279) seedlings (Figure 9B, F). When focused on the repressed DEGs after 8 h of W+FR, 654 and 568 genes were identified in Col-0 and *hy5* seedlings, respectively. A reduced number of genes were repressed in *pif457* (435), whereas the highest number of repressed genes was observed in *hy5 pif457* (690) seedlings (Figure 9C, G). Venn diagrams indicated that, from the total number of DEGs identified in all genotypes (1347 upregulated and 1865 downregulated), a big fraction appeared as upregulated (65.9%: 416 in Col-0, 144 in *hy5*, 159 in *pif457* and 168 in *hy5 pif457*) or downregulated (78.4%: 340 in Col-0, 319 in *hy5*, 282 in *pif457* and 521 in *hy5 pif457*) only in one genotype, whereas the rest appeared in at least two genotypes (Figure 9F-G). These results indicate that after 8 h of simulated shade, the expression of a substantial amount of DEGs does not require PIF457 activity, in contrast to what happens at 1 h.

Regarding the functional prediction, the DEGs belonged to similar GO terms categories in all genotypes (except in *pif457* and *hy5 pif457* after 1 h of W+FR, in which no GO term enrichment was found because of the massive drop in DEG number) (Supplementary Table S5). More importantly, no obvious and specific processes were differentially affected by HY5 (at 1 h) or HY5 and PIF457 at later times that could easily explain the differences in growth detected among the genotypes (Figure 5H). For that reason, we paid attention to the evolution with time of the enrichment of 33 GO terms of biological processes specifically related with shade avoidance, light and growth, and five hormone groups (auxins, gibberellins, ethylene, brassinosteroids and cytokinins) (Table 1) that are known to influence the shade-induced hypocotyl elongation [42, 58–61]. Although most of the enriched GO terms were the same in the various genotypes (Supplementary Tables S5, S6), their p-value changed, reflecting differences in the relative number of genes that participate in a biological process in each genotype (the lower the p-value, the higher the number of DEGs in response to simulated shade involved in a biological process). Regarding the functional predictions of rapidly (1 h) upregulated genes, 12 GO terms related to light, growth and shade avoidance were found as significantly enriched in Col-0 seedlings. Eleven of them were also found in *hy5* with slightly higher p-values (less DEGs contributing to the GO terms) (Table 1). In the case of functional predictions related with hormones, 13 GO terms involved with auxins, gibberellins, ethylene, brassinosteroids and cytokinins were found as significantly enriched in Col-0. As before, most of them (9) were also found in *hy5* with similar or slightly higher p-values as in wild-type seedlings (Table 1). Altogether, only 5 GO terms (response to light intensity, auxin transport, auxin polar transport, response to ethylene and response to cytokinins) were found in Col-0 but not in *hy5*. These results indicate that all these processes (shade avoidance, light and growth, and five hormone groups) depend on the presence of PIF457 to be activated after 1 h of shade treatment.

In the upregulated genes after 8 h of W+FR treatment, from the total of GO terms considered, most were enriched in Col-0 (28), *hy5* (24) and even in the mutant *pif457* (18). A few of them (9) were also enriched in *hy5 pif457* (Table 1). In comparison to Col-0 and *hy5,* p-values in *pif457* and *hy5 pif457* were higher in several GO terms (i.e., relatively less induced genes contributed to enrich in a GO term), such as response to light stimulus, response to red of far-red light and response to auxin stimulus (Table 1). Importantly, several of the biological processes not found in *pif457* after just 1 h of W+FR were identified in the mutant after 8 h and with similar p-values to those of Col-0 and *hy5*. These results indicated that although (1) PIF457 are still playing an important role in regulating several processes after 8 h of W+FR treatment, (2) the early (1 h) leading role of PIF457 after W+FR exposure dissipates after longer periods (8 h) of simulated shade. An exception for this observation was the enrichment in GO terms related with gibberellins, that was absent in *pif457* and *hy5 pif457* after 1 h and 8 h of simulated shade. A different scenario was found in the case of ethylene-related GO terms after 8 h of W+FR, where “ethylene biosynthetic process” and “ethylene metabolic process” were not present in any of the mutants. However, processes related with “ethylene signaling pathway” and “response to ethylene” were enriched in *pif457* and *hy5 pif457* with even slightly lower p-values than in Col-0 (Table 1). Regarding the functional predictions of downregulated DEGs, we found enrichment in a small number of processes related with light and hormones. As for functional predictions of up-regulated DEGs, the few processes enriched after 1 h of W+FR in Col-0 were also enriched in *hy5* but not in the *pif457* or *hy5 pif457* mutants. Overall, there are not many enriched GO terms in the selected processes, particularly after 8 h (Table 1). Together, we concluded that PIF457 and HY5 have a very strong impact in the early shade-regulated changes in gene expression. By contrast, after 8 h of W+FR treatment, the transcriptional responses diverged among genotypes and the influence of PIF457 and HY5, although important, became less apparent.

## DISCUSSION

In the regulation of the *A. thaliana* shade-induced hypocotyl elongation, the function of phyB, PIFs, HFR1 and its dependency on auxin biosynthesis via YUCCA and SAV3 has been well established [7, 11, 12, 18, 31, 49] (Figure 1). The observed genetic interactions between *sav3* and *hfr1*, and *pif7*/*pif457* and *hfr1* (Figure 5A-C) are consistent with this scenario and point to the existence of a branch or module (formed by PIF457-HFR1) directly involved in the rapid and temporal synthesis of IAA observed after exposure to simulated shade (1-4 h), that is dependent on SAV3 and YUC enzymatic activities [12, 49, 58]. Consistently, the enhanced elongation shown by *hfr1* seedlings in response to shade is fully abolished by the auxin biosynthesis inhibitor L-kyn (Figure 2A). We did not find differences on the rapid shade-induced production of auxin in *hfr1* compared with Col-0 (Figure 2B), in contrast with published information [23, 42]. Differences in the light conditions and/or the plant material harvested might account for the lack of impact of HFR1 in the well-established PIF457 role in the promotion of auxin synthesis. This PIF457-HFR1 consensus module appears to be working in controlling SAS responses in seedlings along the whole period of time we have analyzed (2-7 days from germination) (Figure 6) and even in other organs and stages of development, such as petiole length and lamina size in leaves [23].

How the action of phyA and HY5, the other negative regulators analyzed in here, integrates in the consensual SAS regulatory network, is poorly understood (Figure 1). Our genetic analyses with SAS negative and positive regulators (Figures 3-5) hinted that phyA and HY5 act in a different branch than PIF457-HFR1 and SAV3. The enhanced resistance to the auxin biosynthesis inhibitor L-kyn and the auxin polar transport NPA shown by *phyA* and *hy5* seedlings in response to shade, and the attenuated shade-induced rapid IAA production (relative to Col-0) compared to the distinct response of the *hfr1* seedlings further supported this possibility (Figure 2B). The strong elongation in W+FR observed in *hy5 pif457*, *hy5 sav3* and particularly *phyA pif457* and *phyA sav3* hypocotyls (Figure 5D-I) supported that these genotypes elongated even without the shade-induced production of IAA dependent on SAV3 and YUC activities. In the absence of phyA and PIF457, PIF1 and PIF3 might accumulate under shade, bind the DNA and promote the expression of *YUC* genes that would result in auxin biosynthesis and consequently on the elongation of *phyA pif457* hypocotyls [phyA also interacts interact with PIF1 and PIF3 through their Active Phytochrome A-binding motif (APA, needed for active phyA-specific binding) suppressing their regulatory activity [62, 63]. However, this is an unlikely possibility as *phyA sav3* and *hy5 sav3* still are able to elongate. Alternatively, these mutants might elongate either using IAA generated from another biosynthesis pathway where PIF457 and SAV3 are not required or without the need of *de novo* synthesis of IAA. Indeed, it has been shown that IAA can be produced from IAA-conjugated with amino acids molecules in the hypocotyl, IAA that is able to elicit the shade-induced hypocotyl elongation independently of the SAV3-mediated IAA biosynthesis in cotyledons [25]. This increase in IAA potentially produced in the hypocotyl by this alternative pathway does not seem to be high enough for being detected when quantifying IAA in whole *phyA* or *hy5* seedlings (Figure 2B). It seems therefore that the regulatory activity of phyA and HY5 could form a module somehow separated from that of PIF457-HFR1 and the IAA synthetized via the SAV3-YUC pathway.

The participation of HY5 and phyA in the same module is supported by other observations: (1) HY5 is involved in the phyA-mediated down-regulation of early-induced shade avoidance genes in prolonged low R:FR [18], and (2) our growth rate analyses indicate that both phyA and HY5 act very early in the seedling development (days 2-5) (Figure 6). Together, this evidence points to these two factors as having a central importance in repressing the shade-induced growth in this period caused by the inactivation of phyB, providing a temporal framework to sustain the functional separation of these two signaling modules. This phyA-HY5 early suppression action seems fundamental for seedlings establishment and survival soon after germination has occurred in deep shade environments [9], likely by avoiding that the emerging *A. thaliana* seedling would use most of the seed resources in elongating, when overgrowing a dense vegetation canopy is extremely unlikely to succeed. Because deep canopy conditions are usually accompanied by reductions in the light intensity, the mechanisms of elongation are more dependent on dealing with an increased auxin sensitivity [42] that can be modulated directly by phyA action on the stability of the auxin signaling repressors Aux/IAA [10] and by HY5 on the promotion of the expression of negative regulators of auxin signaling [46].

Perception of the R:FR by phyB in the control of the hypocotyl elongation occurs mainly in the cotyledons, where the PIF457-HFR1 module orchestrates a complex transcriptional cascade involving direct activation of *YUC* genes that results in IAA production [12, 40, 43, 64]. This signal is transmitted to the cells of a different organ, the hypocotyl (a typical case of intercellular signaling) [65], where cell elongation, but not division, is promoted [51]. As a consequence, the activity of these module members, such as HFR1, is almost fully abolished by the action of the auxin polar transport inhibitor NPA (Figure 2C). Our cellular analyses indicated that exposure to shade results in a skewed asymmetric bell-shaped growth distribution in which cells 7-8 experienced the maximum elongation in the hypocotyl (Figure 7). Previous observations in various species are consistent with an asymmetric distribution of elongation within stems (but differing in the position of elongation peak) in response to a range of light conditions, such as *A. thaliana* hypocotyls exposed to dark, W [51] or low intensity of blue [43], *Brassica rapa* hypocotyls exposed to W+FR [40] and cowpea epicotyls exposed to end-of-day-FR treatments [66]. However, our work goes further in assigning a role to specific SAS components in the control of growth at the cellular resolution.

Auxin-deficient mutant *sav3* (and *phyA sav3*) limits the shade-induced extra elongation of the hypocotyl cells, indicating that SAV3 (known to act in cotyledons) [49] and transported auxins are needed for the elongation of hypocotyl cells in the lower half of the hypocotyl that occurs in wild-type (and *phyA*) seedlings in W+FR (Figure 7A). The mild redistribution of cell growth observed in the *hfr1* hypocotyls (towards upper cells) (Figure 7B) suggested that the excess of IAA arriving from the cotyledons slightly affected the elongation distribution rather than promoted more growth of the same cells. By contrast, the enhanced elongation observed in *hy5* and *phyA* hypocotyls is caused by the extra cell elongation of the upper part of the hypocotyl, a growth redistribution that happens even in the absence of IAA arriving from the cotyledons (as observed in *phyA sav3* hypocotyls) (Figure 7A, C). These analyses spatially help to distinguish the regulatory activity of the module in which HFR1 participates, that acts in the middle-lower half of the hypocotyl and it is highly SAV3-dependent, from that of phyA-HY5, that mainly represses cell elongation in the upper half. Together, the spatial and temporal framework provided by our results suggest that, in a wild-type background, the strong repression imposed by phyA-HY5 in the upper half of the hypocotyls at the beginning of seedling development progressively dissipates and the elongation of the upper cells in the hypocotyl takes place. This temporal and spatial separation of the PIF457-HFR1 (together with SAV3) and phyA-HY5 regulatory activities would be consistent with an acropetal gradient of hypocotyl growth (from the base to the top) in response to simulated shade, as it was observed in both dark- and W-grown seedlings [51].

An additional level of regulation refers to when the different SAS components and modules act relative to the beginning of the simulated shade exposure, as well as their level of molecular interaction. Our expression analyses indicate that PIF457 are essential to modulate gene expression immediately after shade exposure. In contrast with *PIF457*, *HY5*-deficiency enhanced the number of DEGs after 1 h (1142) compared to Col-0 (563). Importantly, the vast majority of these rapid DEGs found in the *hy5* background also required PIF457 to be shade-regulated (20 DEGs in *hy5 pif457*) (Figures 8, 9D-E) indicating that *PIF4*, *PIF5* and *PIF7* are epistatic over *HY5* at this early time after shade exposure. The observed epistasis suggests that the transcriptional activity of these two factors might integrate and connect the two mentioned branches at these initial stages after shade exposure. Previously, it has been demonstrated physical interaction between HY5 and PIFs or convergence of their transcriptional activities in non-shade-related processes [52, 54, 67]. Thus, PIFs and HY5 could be key players connecting the two regulatory modules rapidly after shade exposure.

The crosstalk between PIF457 and HY5 activities, however, seems dynamic and changes with longer times of shade exposure. After 8 h, the transcriptome was clearly affected by shade even in the absence of PIF457 (Figure 9B-C, F-G), reflecting that an important percentage of the expression changes caused after shade perception by phyB happens bypassing PIF457 activity. Hence, expression of these DEGs depends either on other PIFs (e.g., PIF1 and PIF3) or on the effect exerted by unknown but non-PIF regulators whose transcriptional activity is also connected to the reduction in phyB activity under W+FR. In addition, after 8 h of shade exposure, the transcriptome divergence between the various genotypes, even between *pif457* and *hy5 pif457* (Figure 9F-G), point to a change in the molecular relationship of *PIFs* and *HY5* that appear in this moment to act independently from each other. What sustains this dynamic relationship is unknown, although it might involve changes in the accessibility of these regulators to the same target promoters with time triggered by shade perception (e.g., caused by the increase in the abundance of transcriptional regulators-cofactors that can affect their DNA-binding abilities), a shade-induced divergence of their spatial pattern of expression that impedes PIFs and HY5 to be expressed in the same cells, and/or by epigenetic processes that alter chromatin compaction, also known to influence the accessibility and binding of transcription factors to regulatory elements in the DNA [39].

Apart from the dramatic reduction in the number of DEGs in the *pif457* and *hy5 pif457* samples harvested after 1 h of simulated shade, no process (based on GO terms) was clearly associated with HY5 activity at 1 h, or even with PIF457 or HY5 after 8 h of simulated shade exposure. However, the activities of these components (and the modules they represent) are not totally independent, and several processes are co-regulated by both HY5 and PIFs such as light, growth and auxin related processes, suggesting that some aspect of PIFs and HY5 transcriptional activities converge within 8 h of shade exposure. However, the observed transcriptomic changes, that do not associate with the growth response, could not explain the almost wild-type elongation of *hy5 pif457* hypocotyls (Figure 5).

In the SAS regulation, PIFs are usually presented as positive regulators by promoting the expression of genes involved in hypocotyl elongation. Our RNA-seq analyses support that they also have an important function in the repression of gene expression, as it has been previously described for some PIFs in shade-induced processes related with metabolic or architectural responses (hence, not-related with cell elongation) [68–70]. Similarly, although HY5 acts mainly inducing gene expression [71], it has an important role in the shade-repression of genes that, after just 1 h of shade exposure, requires PIF457 (Figure 9D-E).

Our findings propose a model for the regulation of shade-induced hypocotyl elongation that incorporates the temporal and spatial functional importance of the various SAS regulators analyzed in here. These components are grouped in two main modules or branches: (1) a well-defined pathway in which PIF457-HFR1 participate, it is highly dependent on auxin produced via SAV3 and YUCs mostly in the cotyledons, acts along all seedling development (from day 2 to 7 from germination, although apparently more strongly at the end of the period analyzed) and targets cells in the middle-lower region of the hypocotyl; and (2) a less well-characterized pathway with phyA and HY5 as main components, that is less dependent on SAV3-related auxin biosynthesis and polar transport, it has an important role in the early seedlings development (day 2 to 5 after germination) and targets cells in the upper region of the hypocotyl. Our analyses also show that in these processes, PIF457 transcriptional activity is fundamental at 1 h of W+FR and its importance dissipates at later times (8 h). By contrast, importance of HY5 regulatory role increases at longer times of shade exposure, when its expression is also reported to enhance [18], likely because of the delayed accumulation of phyA. Importantly, the molecular interaction between these transcriptional regulators is dynamic and moves from epistasis, soon after shade exposure, to additivity, at later hours (based on both transcriptomic and hypocotyl elongation experiments).

### MATERIAL and METHODS

### Plant material and growth conditions

All the *Arabidopsis thaliana* plant material used was in the Columbia-0 (Col-0) background. Mutants used in this study were described before: *phyA-501* [4], *hy5-2* [35]*, hfr1-5 (*Roig-Villanova etal_2007 [14], *pif7-1* [11] and *sav3-5*, also known as *wei8-4* / *tir2-3* [72]. The multiple mutants *pif457* (*pif4-101 pif5-3 pif7-1*) [23], *phyA-211 hfr1-101* [73] and *phyA-211 phyB-*9 [74] used in this study were described elsewhere. To produce seeds of the various *A. thaliana* genotypes, plants were grown in the greenhouse under long day photoperiod (16 h light, 8 h dark).

Fluence rates were measured with a Spectrosense2 meter associated with a 4-channel sensor (SkyeInstruments Ltd., www.skyeinstruments.com), which measures PAR (400–700 nm) and 10 nm windows in the blue (464 – 473 nm), R (664 – 674 nm) and FR (725 – 735 nm) regions.

### Pharmacological treatments

When indicated, the medium was supplemented with different concentrations of L-kynurenine (L-kyn, Sigma-Aldrich) or NPA (Duchefa). L-kyn was dissolved at 50 mM in DMSO. NPA was dissolved at 5 mM in DMSO. Stock solutions were kept at -20°C until use.

### Genetic crosses and genotyping

Mutants were crossed to generate the following multiple mutants: *phyA hy5* (*phyA-501 hy5-2*), *phyA pif7* (*phyA-501 pif7-1*), *phyA hfr1* (*phyA-501 hfr1-5*), *phyA sav3 (phyA-501 sav3-5)*, *hy5 pif7* (*hy5-2 pif7-1*), *hy5 hfr1* (*hy5-2 hfr1-5*), *hy5 sav3* (*hy5-2 sav3-5*), *hfr1 pif7* (*hfr1-5 pif7-1)*, *hfr1 pif457* (*hfr1-5 pif457*), *hy5 pif457* (*hy5-2 pif457*) and *phyA pif457* (*phyA-501 pif457*). After crosses, seedlings in the segregating F2 generation were pre-selected searching by the predicted phenotypes, if any. In any case, the genetic identity of the plants was established by genotyping the pre-selected plants by PCR using specific primers (Supplementary Table S7).

### Measurements of hypocotyl length

For hypocotyl growth assays, seeds were sterilized and sown in solid agar plates without sucrose (GM–; 0.215% (w/v) MS salts plus vitamins, 0.025% (w/v) MES pH 5.80) [75]. After 3-6 days of stratification, plates were incubated in growth chambers at 22°C under continuous white light (W) provided by 4 cool-white vertical fluorescence tubes for 2 days (PAR of 20–25 umol·m^-2^·s^-1^, R:FR of about 1.58). After that time, plates were either maintained in W or transferred to simulated shade (W+FR) for 5 days. Simulated shade was generated by enriching W with supplementary FR provided by 4 horizontal LED lamps (PAR of 20–25 umol·m^-2^·s^-1^, R:FR of about 0.02). Details of the resulting light spectra have been described before [8]. At day 7, seedlings were flattened down on the petri dishes and pictures of them were taken. Each biological replicate corresponded to ∼25 seedlings per treatment and genotype. Experiments were done with at least 3 biological replicates. Hypocotyl measurements were carried out by using the National Institutes of Health (NHS) ImageJ software (Bethesda, MD, USA; http://rsb.info.nih.gov/). Hypocotyl measurements from the different biological replicates were averaged.

### Hypocotyl measurements for the temporal analyses

Seedlings were grown for up to 7 days either in W or W+FR, as described in the previous section. In these experiments, hypocotyl length measurements were made daily from pictures taken from plants of different ages, from day 2 until day 7 after germination (6 time points). By subtracting hypocotyl length of two consecutive days, growth rate (mm·day^-1^) from day 2 to day 6 was calculated for each genotype and light treatment (W and W+FR).

Each biological replicate corresponded to ∼25 seedlings per treatment, genotype and time point. Experiments were done with 3 biological replicates. Hypocotyl measurements from the different biological replicates were averaged. These averaged data were used to calculate the growth rate.

### Hormone analyses

About 50 seedlings per biological replica of the different genotypes and treatments (that ranged from 80-120 mg) were immediately frozen in liquid nitrogen. Hormone extraction and analysis were performed as described [76] with a few modifications. Briefly, around 100 mg of fresh material was extracted in 1 mL of 50% acetonitrile (v/v) prepared with ultrapure water, adding 2.5 ng of [^2^H_5_]IAA as internal standard in a ball mill (MillMix 20, Domel) for 10 min at 17 rps, followed by 5 min of sonication. After sonication, the samples were centrifuged at 4000g at 4°C for 10 min. Finally, supernatants were filtered through SPE columns (OASIS HLB 30 mg 1 cc, Waters), recovering the eluent. Finally, 0.5 mL of 30% acetonitrile (v/v) prepared in ultrapure water was added to the SPE columns and the eluent was recovered joint to the previous ones.

Chromatographic separations were performed on a reverse-phase C18 column (gravity, 50 3 2.1 mm, 1.8-mm particle size, Luna-Omega, Phenomenex) using a acetonitrile:water (both supplemented with 0.1% formic acid) gradient at a flow rate of 300 mL/min. IAA was detected with a triple quadrupole mass spectrometer (Micromass) connected online to the output of the column though an orthogonal Z-spray electrospray ion source. Finally, IAA content was quantified by interpolation in a standard curve prepared with commercial IAA (Sigma) using the Masslynx v4.2 software.

### Cell length measurements along the hypocotyl axis for spatial analysis

For the cell length measurements, about 100 seedlings were grown either in W or W+FR, as previously described. On day 7, hypocotyls were measured and the mean value for each group (genotype and treatment) was calculated. About 15 individuals with a hypocotyl length of the estimated averaged value ± 5 % were selected. Cotyledons and roots were removed from these seedlings and the remaining hypocotyls fixed and stained with Calcofluor White (Sigma-Aldrich) to visualize cell walls. Briefly, hypocotyls were submerged in a 1x PBS solution [137 mM NaCl (8.06 g/L), 2.7 mM KCl (0.22 g/L), 10 mM Na_2_HPO_4_ (1.15 g/L), 18 mM KH_2_HPO_4_ (0.20 g/L)] with 4% (w/v) paraformaldehyde (PFA) for 60 min at room temperature. Then, hypocotyls were washed twice for 1 min in 1x PBS and cleared after transferring them to ClearSee solution [10% (w/v) Xylitol (Sigma), 15% (w/v) Sodium deoxycholate (Sigma), 25% (w/v) Urea (Sigma)]. The clearing was carried out for at least 1 week at room temperature. Before taking images, hypocotyls were stained with 100 µg·ml^-1^ Calcofluor in ClearSee solution for 120 min, and washed twice with ClearSee solution for 2 days [77].

Images of fixed and stained plant material were taken by using confocal microscopy (Zeiss LSM 780). Calcofluor White stained samples were imaged with 405 nm excitation and detected at 425-475 nm [78]. Cell growth measurements were carried out using the NHS ImageJ software on the obtained pictures. At least 15 cells of 2 cell files per hypocotyl from 7 seedlings were measured for each genotype and growth condition.

### RNA extraction and gene expression analyses

Seven-day old seedlings grown in W or W+FR were harvested (about 35 mg per sample) and frozen in liquid nitrogen. RNA was extracted using commercial kits (Maxwell® RSC Plant RNA kits; www.promega.com) and quantified using *NanoDrop^TM^ 8000 spectrophotometer* (ThermoFischer Scientific^TM^). Two µg of total RNA were retrotranscribed to cDNA in a final volume of 20 µL by using the Transcriptor First Strand cDNA synthesis KIT (Roche, www.roche.com) or the NZY First-strand cDNA synthesis kit, separate oligos (NZYtech) following the manufacturer’s instructions. Subsequently cDNA was diluted ten-fold and stored at -20°C for further analysis.

Relative mRNA abundance was determined via Real-Time Quantitative Polymerase Chain Reaction (RT-qPCR) in a final volume of 10 µL made up of 0.3 µM of both, forward and reverse primers, 5 µL of the LightCycler 480 SYBR Green I Master Mix (Roche) and 2 µL of ten-fold diluted cDNA [8]. The RT-qPCR was carried out in *LightCycler 480 Real-Time PCR system* (Roche). The analysis was performed with three independent biological replicates (∼30 seedlings per biological replicate) for each condition and three technical replicates for each biological replicate. *ELONGATION FACTOR 1α* (*EF1α*) was used as endogenous reference genes to normalize the expression of the genes of interest. Primers used for the RT-qPCR analyses are provided in Supplementary Table S8.

### Statistical analyses

These analyses were carried out using the Real Statistics Resource Pack, an Excel add-in that extends Excel’s standard statistics capabilities. For the statistical analyses, we compared three values corresponding to three replicates in the case of relative expression and hypocotyl length.

### RNA-sequencing

Total RNA for sequencing was obtained as in the expression analyses by RT-qPCR. Library preparation was performed from three biological replicates and sequenced at the Centre Nacional d’Anàlisi Genòmica (CNAG-CRG) on an Illumina NovaSeq6000 in paired-end 50 bp read length. Mapping by TAIR10 genome and Limma comparisons were performed to obtain the False Discovery Rate (FDR) and the Fold Change (FC) for each gene. Differentially expressed genes (DEGs) were considered when FDR P-value was <0.05 and FC > |1.5|.

### GO enrichment

The list of DEGs (Supplementary Tables S1-S4) were used to identified the enrichment in GO terms using the agriGO online analyses tool.

## ACKNOWLEDGEMENTS

We are grateful to Christian Fankhauser (University of Lausanne, Switzerland) for providing *pif457* and *phyA-211 hfr1-101* seeds; to Pablo Cerdán (Fundación Instituto Leloir, Argentina) for providing *phyA-211 phyB-9* seeds; and to Javier Brumós (IBMCP) for comments on the manuscript. PP-A was funded by an EMBO ShortTerm fellowship that covered his stay at the ETH, Zurich. MU-B is supported by a predoctoral fellowship from Spanish Ministerio de Universidades - FPU20/05486), JM-R has received funding from the European Union’s Horizon 2020 research and innovation program under the Marie Slodowska-Curie (H2020-MSCA-IF-2017) grant agreement 797473. Our research is supported by grants BIO2017-85316-R from MCIN/AEI/10.13039/501100011033, “ERDF A way of making Europe”, PID2020-115782GB-I00 from MCIN/AEI/10.13039/501100011033, PROMETEU/2021/065 from Generalitat Valenciana and the EC-H2020-PRIMA UToPIQ project funded by AEI (PCI2021-121941). We also acknowledge the support of the CERCA Programme / Generalitat de Catalunya.

## SUPPLEMENTARY DATA

**Table 1.** Summary of significance of the enrichment of a subset of selected GO terms in Col-0 (wild-type), *hy5*, *pif457* and *hy5 pif457* seedlings. GO term enrichments were obtained using the agriGO online analyses tool and the up-(left section) and down-regulated (right section) DEGs summarized in Supplementary Tables S1-S4. Groups presented were selected because p-values are significantly overrepresented in at least one of the four genotypes with W+FR for 1 and/or 8 h. Cell color indicates the significance of the enrichment of the GO term (green color, p-value < 0.05; light orange color, p-value > 0.05).

**Supplementary Figure S1.** (Supports Figure 4). Hypocotyl length in W and W+FR of Col-0 **(A)**, *phyA-211*, *hfr1-5, phyA-211 hfr1-101*, **(B)** *hy5-2*, *hfr1-5*, *hy5-2 hfr1-1*, **(C)** *phyA-501, hy5-2* and *phyA-501 hy5-2.* Seedlings were germinated and grown as indicated in Figure 3A. Values are means and SE of three independent samples. Different letters denote significant differences (one-way ANOVA with the Tukey test, P-value < 0.05) among means.

**Supplementary Figure S2.** (Supports Figure 5). Hypocotyl length in W and W+FR of Col-0 **(A)** *hfr1-5*, *pif7-1*, *hfr1-5 pif7-1*, **(B)** *hfr1-5*, *pif457*, *hfr1-5 pif457*, **(C)** *hfr1-5*, *sav3-5*, *hfr1-5 sav3-5*, **(D)** *phyA-501*, *pif7-1*, *phyA-501 pif7-1*, **(E)** *phyA-501*, *pif457*, *phyA-501 pif457*, **(F)** *phyA-501*, *sav3-5*, *phyA-501 sav3-5*, **(G)** *hy5-2*, *pif7-1*, *hy5-2 pif7-1*, **(H)** *hy5-2*, *pif457*, *hy5-2 pif457*, and **(I)** *hy5-2*, *sav3-5*, *hy5-2 sav3-5.* Seedlings were germinated and grown as indicated in Figure 3A. Data of section **A** (Col-0, *hfr1-5*, *pif7-1*, *hfr1-5 pif7-1*) have been already presented (Figure 6B in [31]). Values are means and SE of three independent samples. Different letters denote significant differences (one-way ANOVA with the Tukey test, P-value < 0.05) among means.

**Supplementary Figure S3.** (Supports Figure 6). Growth rate (GR) of **(A)** *phyA-501*, *phyB-9*, **(B)** *hfr1-5*, *hy5-2* and *phyA-501* relative to Col-0 grown in the same light conditions. Values were calculated by subtracting to the GR of the mutant those of Col-0 grown in the same light conditions (data presented as Figure 6).

**Supplementary Figure S4.** (Supports Figure 7). Representative confocal images of Col-0 hypocotyls grown under W or W+FR to measure epidermal cell length.

**Supplementary Figure S5.** (Supports Figure 7). Length in W **(A, C, D)** and W+FR **(B, E, F)** for each of the 20 epidermal cells (from bottom to top) that conform a file in hypocotyls of Col-0, (**A, B)** *sav3-5*, *phyA-501*, *phyA-501 sav3-5*, **(C, E)** *hfr1-5* and **(D, F)** *hy5-2*. Seedlings were germinated and grown as indicated in Figure 3A. Red arrows point to the cell that presents a higher difference in length. Values are mean lengths and SE of at least 15 cells of 2 cell files per hypocotyl from 7 different hypocotyls per genotype and growth condition.

**Supplementary Figure S6.** Correlation between peak of cell number (from Figure 7) and difference in length in W+FR and W hypocotyls (HYP_W+FR_ – HYP_W_) (from Figure 5). Tendency line was dne using a polynomic function of 1 degree without including the *phyA sav3* pair.

**Supplementary Table S1.** Bioset of up-regulated DEGs in Col-0 seedlings in response to 1 h of W+FR.

**Supplementary Table S2.** Bioset of up-regulated DEGs in *hy5* seedlings in response to 1 h of W+FR.

**Supplementary Table S3.** Bioset of up-regulated DEGs in pif457 seedlings in response to 1 h of W+FR.

**Supplementary Table S4.** Bioset of up-regulated DEGs in hy5 pif457 seedlings in response to 1 h of W+FR.

**Supplementary Table S5.** Functional enrichment groups based on GO term analyses.

**Supplementary Table S6.** Functional enrichment groups based on GO term analyses.

**Supplementary Table S7.** Primers used for genotyping the different lines.

**Supplementary Table S8.** Primers used for gene expression analyses.

