## Supplementary figures and images for "Temporal and spatial frameworks supporting plant responses to vegetation proximity"

### Supplemental figures

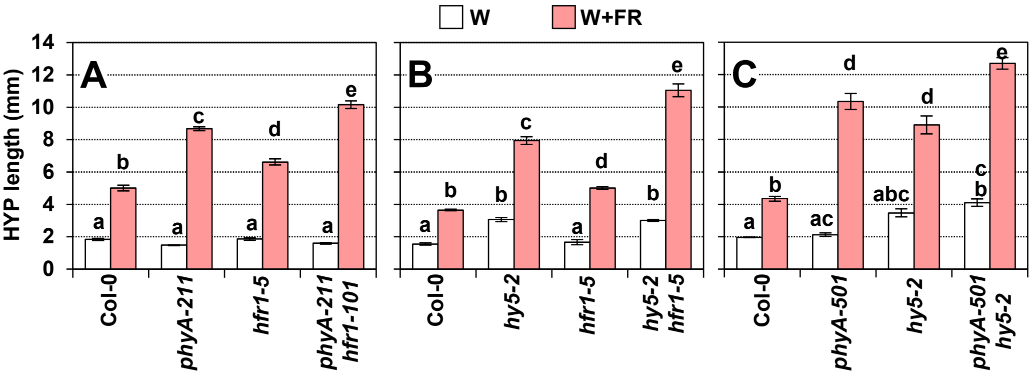

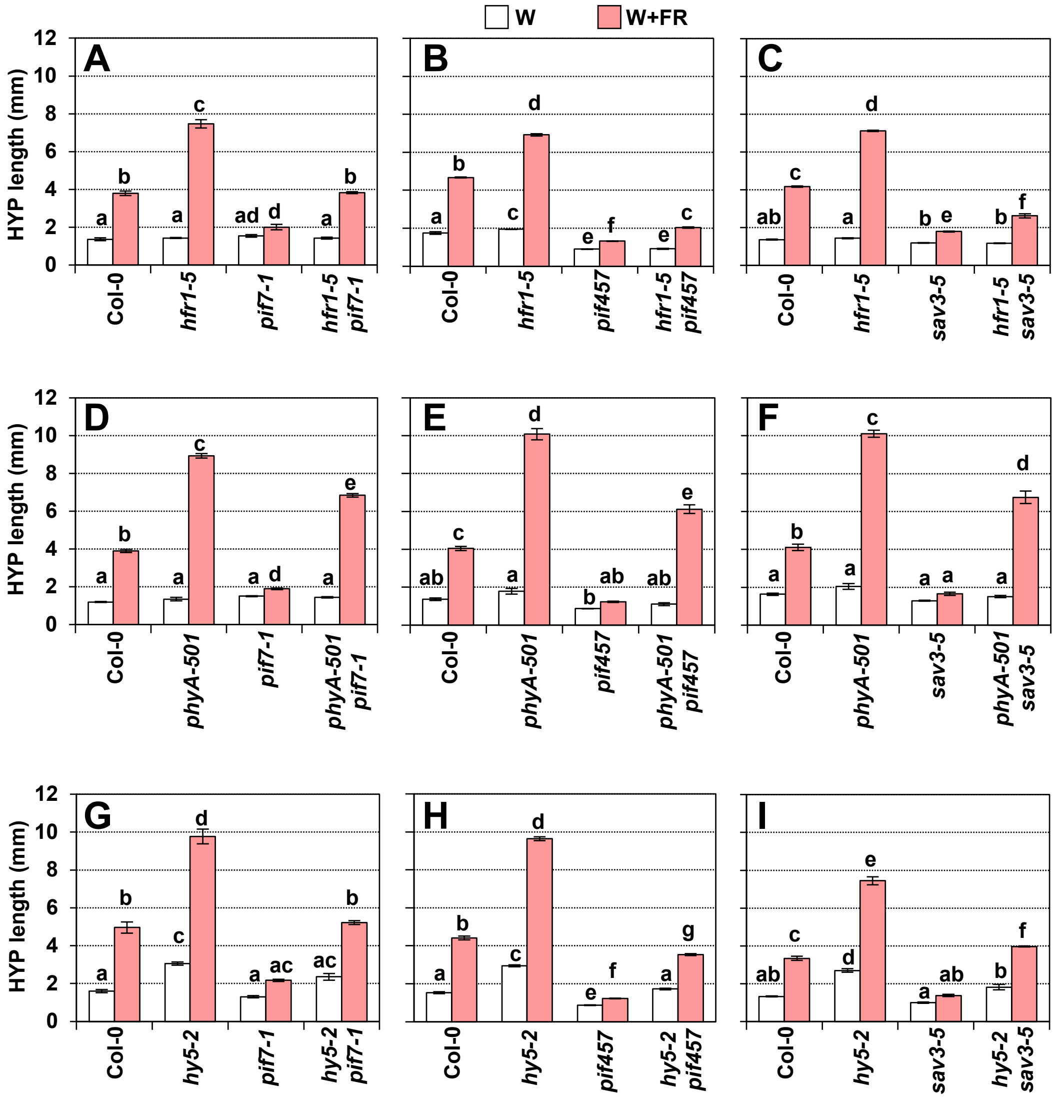

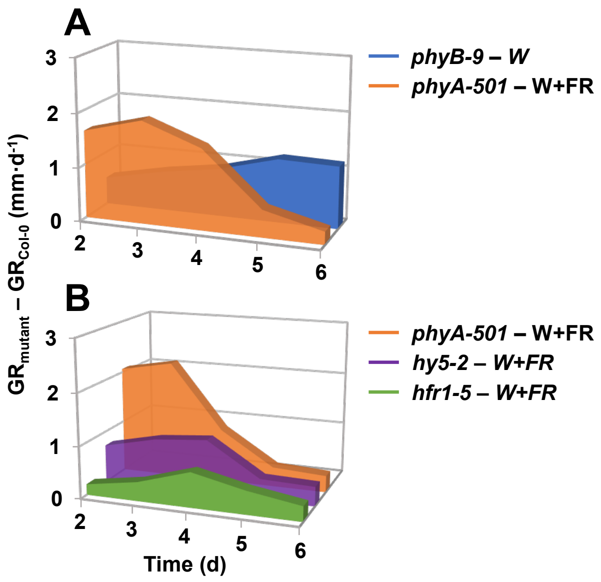

#### Figure S4

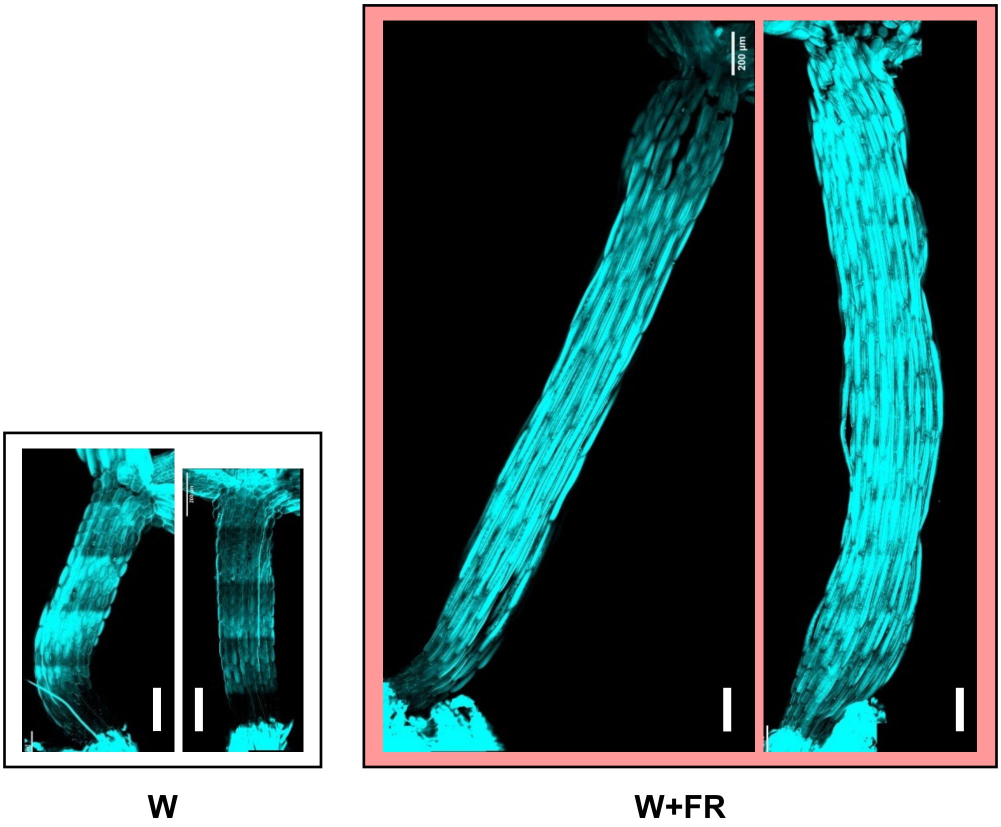

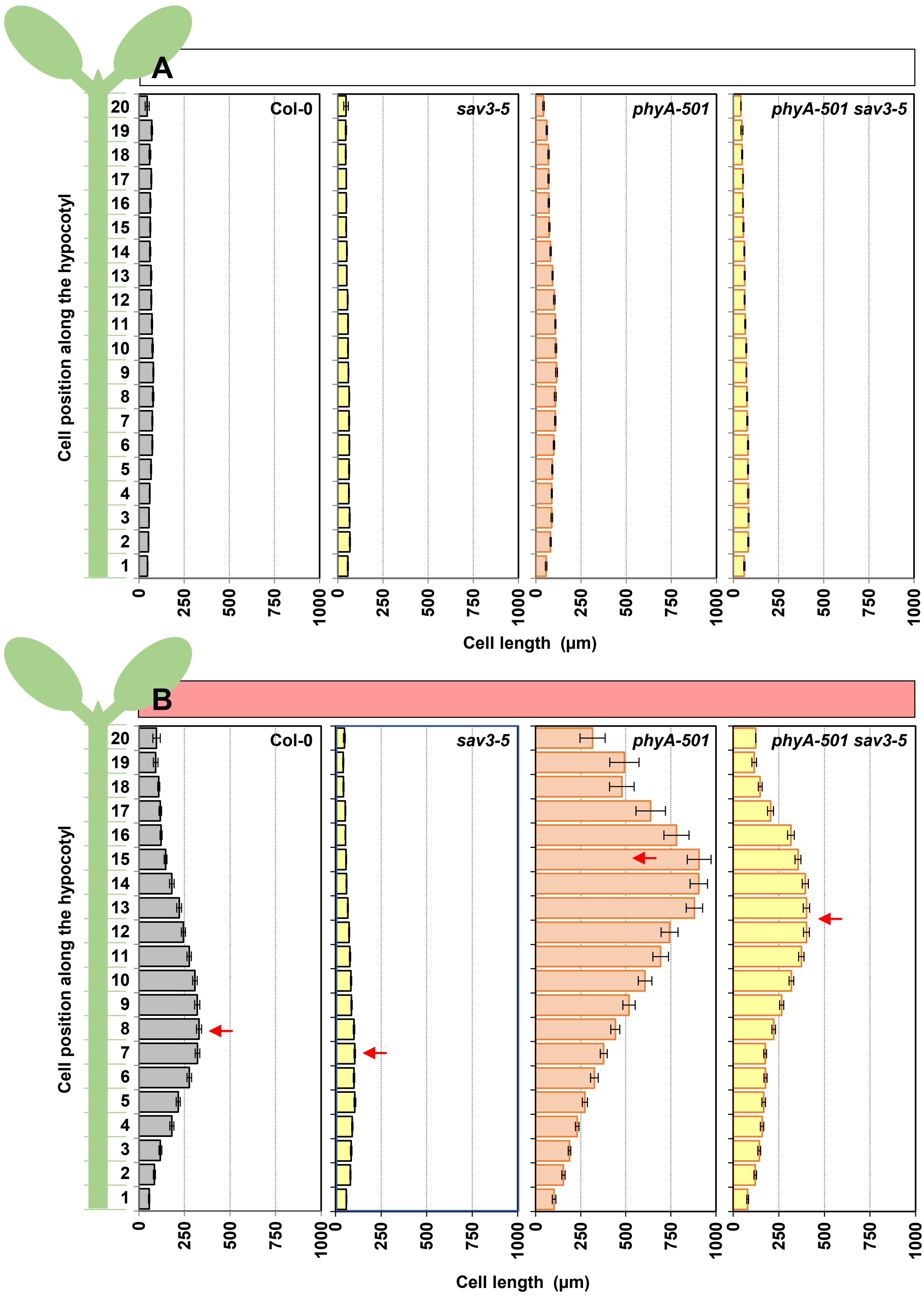

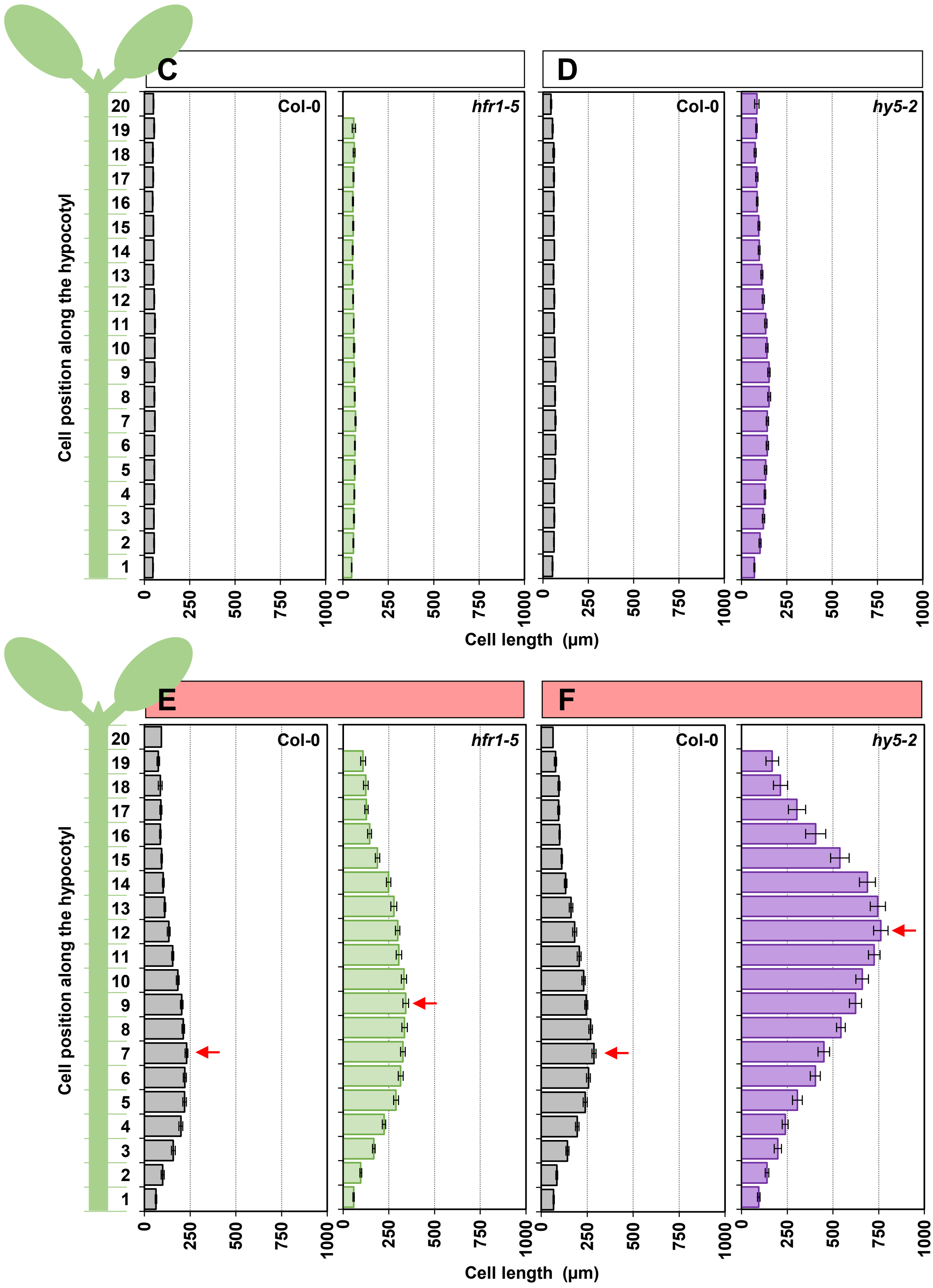

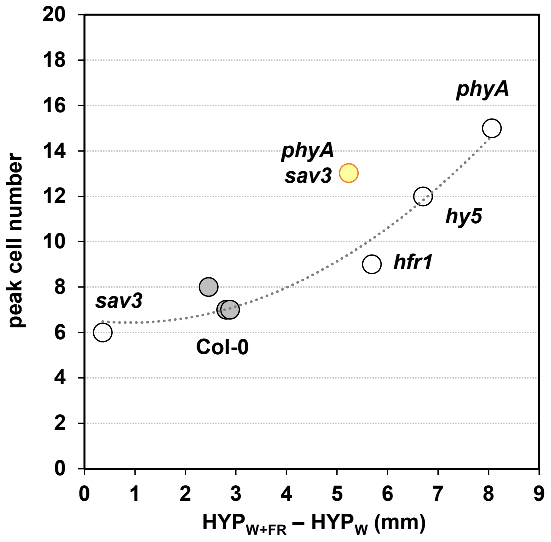
